# RAD18 opposes transcription-associated genome instability through FANCD2 recruitment

**DOI:** 10.1101/2022.06.24.497439

**Authors:** James P. Wells, Emily Yun-chia Chang, Leticia Dinatto, Justin White, Stephanie Ryall, Peter C. Stirling

## Abstract

DNA replication is a vulnerable time for genome stability maintenance. Intrinsic stressors, as well as oncogenic stress, can challenge replication by fostering conflicts with transcription and stabilizing DNA:RNA hybrids. RAD18 is an E3 ubiquitin ligase for PCNA that is involved in coordinating DNA damage tolerance pathways to preserve genome stability during replication. In this study, we show that RAD18 deficient cells have higher levels of transcription-replication conflicts and accumulate DNA:RNA hybrids that induce DNA double strand breaks and replication stress. We find that these effects are driven in part by failure to recruit the Fanconi Anemia protein FANCD2 at difficult to replicate and R-loop prone genomic sites. While RAD18 is important for FANCD2 localization after splicing inhibition or aphidicolin treatment, it is comparatively less important during interstrand crosslink repair. Thus, we highlight a RAD18 dependent pathway promoting FANCD2-mediated suppression of R-loops and transcription-replication conflicts.

**AUTHOR SUMMARY:** Genome instability, a state in which cells acquire mutations more quickly, drives cancer initiation and progression. DNA is normally protected from such mutations by a host of specialized factors that recognize and repair potentially damaging circumstances. Here we describe under-appreciated links between a protein called RAD18, which recognizes potentially damaging stress, and a phenomenon called a transcription-replication conflict. If not repaired such conflicts can lead to DNA breaks. We find that RAD18 plays an important role in recruiting another repair protein called FANCD2 to sites of transcription-replication conflicts. By doing so, RAD18 protects the human genome from damage when it is faced with stresses such as those seen in cancer cells. This work provides new insight to how the FANCD2 protein is brought to potentially damaging sites in our genomes.

## Introduction

RAD18 is a conserved E3 ubiquitin ligase that coordinates multiple DNA repair and damage tolerance pathways including post-replication repair (PRR) pathways, homologous recombination (HR), Fanconi Anemia (FA), and break-induced replication (BIR) (1–5). RAD18 has numerous impacts on genome instability, driven mostly through regulated ubiquitylation of the primary RAD18 substrate, Proliferating Cell Nuclear Antigen (PCNA), which acts as a ring-shaped homotrimer encircling DNA as a sliding clamp component of the replisome (6). PCNA is considered the principal conductor at the replication fork, acting as a scaffold to promote the replication stress response, facilitating chromatin assembly, regulating cell proliferation and apoptosis, and coordinating multiple DNA repair pathways (7).

Ubiquitination of PCNA at lysine 164 by RAD18 initiates a series of choices that, depending on the context and action of downstream factors, trigger DNA damage tolerance. Mono-ubiquitination can serve to activate the translesion synthesis (TLS) pathway to coordinate specialized polymerases that are able to replicate over damaged nucleotides at stalled replication forks (1). However, several of these TLS polymerases are intrinsically error-prone and restart replication at the cost of increased mutations. Extension of mono-ubiquitin into K63-linked polyubiquitination by factors like HLTF-UBC13-MMS2 initiates a recombination-based template switching lesion bypass mechanism (8). The regulation of decisions around TLS, template switching or other pathways has been reviewed elsewhere (9, 10).

The action of RAD18 on PCNA has also been linked to the FA repair pathway, a network of DNA repair genes that coordinate to repair covalent links between DNA called interstrand crosslinks (ICLs). Fanconi Anemia is a genetic disease characterized by bone marrow failure and cancer predisposition and usually caused by loss of function in any of the FA pathway genes (11). This link between RAD18 and PCNA was originally identified through the effect of RAD18 on TLS, which is also an important part of ICL repair by the FA pathway (12). Subsequently, it was shown that loss of RAD18 prevents full ubiquitination and activation of FANCD2 in response to bulky DNA adducts (13) or cisplatin (14) through a mechanism in which FANCL, the E3-ligase for FANCD2, is recruited to RAD18-ubiquitylated PCNA. RAD18 also seems to impact functions of FANCD2 that are independent of ICL repair as it was suggested to act upstream of FANCD2, BRCA2 and RAD51 in the context of HR-repair of camptothecin induced replication fork blockage (15, 16). Relating this to the present work, Topoisomerase I, the target of camptothecin, is also a critical factor opposing transcription-replication conflicts (17).

The canonical FA pathway functions in ICL repair, but has also been implicated in the response to transcription-replication conflicts. Such conflicts can occur in both co-directional and head-on orientations, however head-on conflicts have a more deleterious impact on replication progression, possibly by stabilizing R-loops (18, 19). R-loops are three stranded structures formed upon the reannealing of nascent RNA to the DNA template, that can serve both normal and pathological functions in genome stability (20, 21). FANCA, FANCD2, FANCL, FANCM and FA proteins such as BRCA1 and BRCA2 have all been suggested to mitigate R-loop accumulation and associated DNA damage (22–25). The translocase activity of FANCM has been suggested to mitigate R-loops in general and specifically at ALT-telomeres in association with the BLM-TOP3A-RMI1 complex (26). In addition, the FANCI-FANCD2 complex was suggested to directly bind R-loops and promote FA activation (27). However, we do not fully understand the regulation and function of the FA pathway at transcription-replication conflicts and R-loops. Both ubiquitin-dependent (24) and -independent functions of FANCD2 have been implicated in mitigating the effects of transcription-replication conflicts (28), albeit in different stress contexts.

Recent work on the alternative replication factor C (RFC) subunit ATAD5/Elg1, which unloads PCNA normally and during stresses, has implicated PCNA as a hub of transcription-replication conflict regulation (29). Indeed, our own previous work suggested PCNA in yeast may be an important anti-R-loop mechanism since *elg1*Δ was synthetic lethal with RNaseH-deficiency, and *rad18*Δ caused an increase in DNA:RNA hybrid staining with the S9.6 antibody (30). Here, we set out to follow up on these observations by investigating the influence of RAD18 on transcription-replication conflicts in human cell lines. We show that RAD18 loss results in the accumulation of R-loops in several cell lines but that RAD18 depletion does not further enhance R-loops in PCNA K164R mutant cells. R-loop-dependent replication stress and DNA damage is enhanced in RAD18 deficient cells. Chemical treatments that cause replication stress through R-loop accumulation induced FANCD2 foci, and were significantly dependent on RAD18 status compared to mitomycin C (MMC), which induces ICLs. The recruitment of FANCD2 to common fragile sites (CFS) stimulated by aphidicolin (APH) was significantly reduced in RAD18 deficient cells. Together our study defines the importance of RAD18-mediated PCNA ubiquitination in recruiting proteins of the FA pathway to difficult to replicate regions associated with R-loops and transcription-replication conflicts.

## Materials and Methods

### Cell culture and transfection

Human embryonic kidney (HEK) 293T, mouse embryonic fibroblasts (MEF) and HeLa cells were cultivated in Dulbecco’s modified Eagle’s medium (DMEM) (Stemcell technologies) supplemented with 10% fetal bovine serum (Life Technologies) in 5% CO2 at 37°C. Retinal pigment epithelium (RPE1) cells were cultured as above in DMEM with F-12 modification.

For RNA interference in HeLa, RPE1, HEK 293T, and MEF cells, cells were transfected with siGENOME SMARTpool siRNAs from Dharmacon, which are pools of four siRNA to minimize off-target effects (Non-targeting siRNA Pool #1 as si-Cont, si-RAD18, si-FANCD2, si-FANCL). Transfections were done with Dharmafect1 transfection reagent (Dharmacon) according to manufacturer’s protocol and harvested 48 h after the siRNA administration. For experiments with overexpression of GFP or nuclear-targeting GFP-RNaseH1 (gift from R. Crouch), transfections were performed with Lipofectamine 3000 (Invitrogen) according to manufacturer’s instructions 24 h after the siRNA transfections.

### CRISPR/CAS9 RAD18 KO cell line generation

For RAD18 gene knockout, we utilized the Alt-R™ CRISPR-Cas9 System integrated DNA technologies (IDT). Custom crRNAs was designed using Custom Alt-R® CRISPR-Cas9 guide RNA tool and both crRNAs and tracrRNAs were received from IDT (**Table S1**). RPE-1-hTERT P53^-/-^ cells stably expressing Cas9 nuclease were reverse transfected (RNAiMAX reagent, Thermo Fisher Scientific) with the Alt-R CRISPR-Cas9-crRNA complexed with ALt-R CIRPSR-Cas9 tracrRNA - ATTO 550 (final concentration of 10 nM). FACS cell sorting isolated ATTO 550-positive cells and cells were expanded and sorted again into single cell populations on a 96-well plate. Knockout scores were determined for clones and selected using the synthego ICE tool. RAD18 knockout positive cells were confirmed with PCR using Q5 High Fidelity Master Mix (NEB), sanger sequencing, and western blot (**Table S1**)(**Figure 1, Figure S1)**.

**Figure 1.**
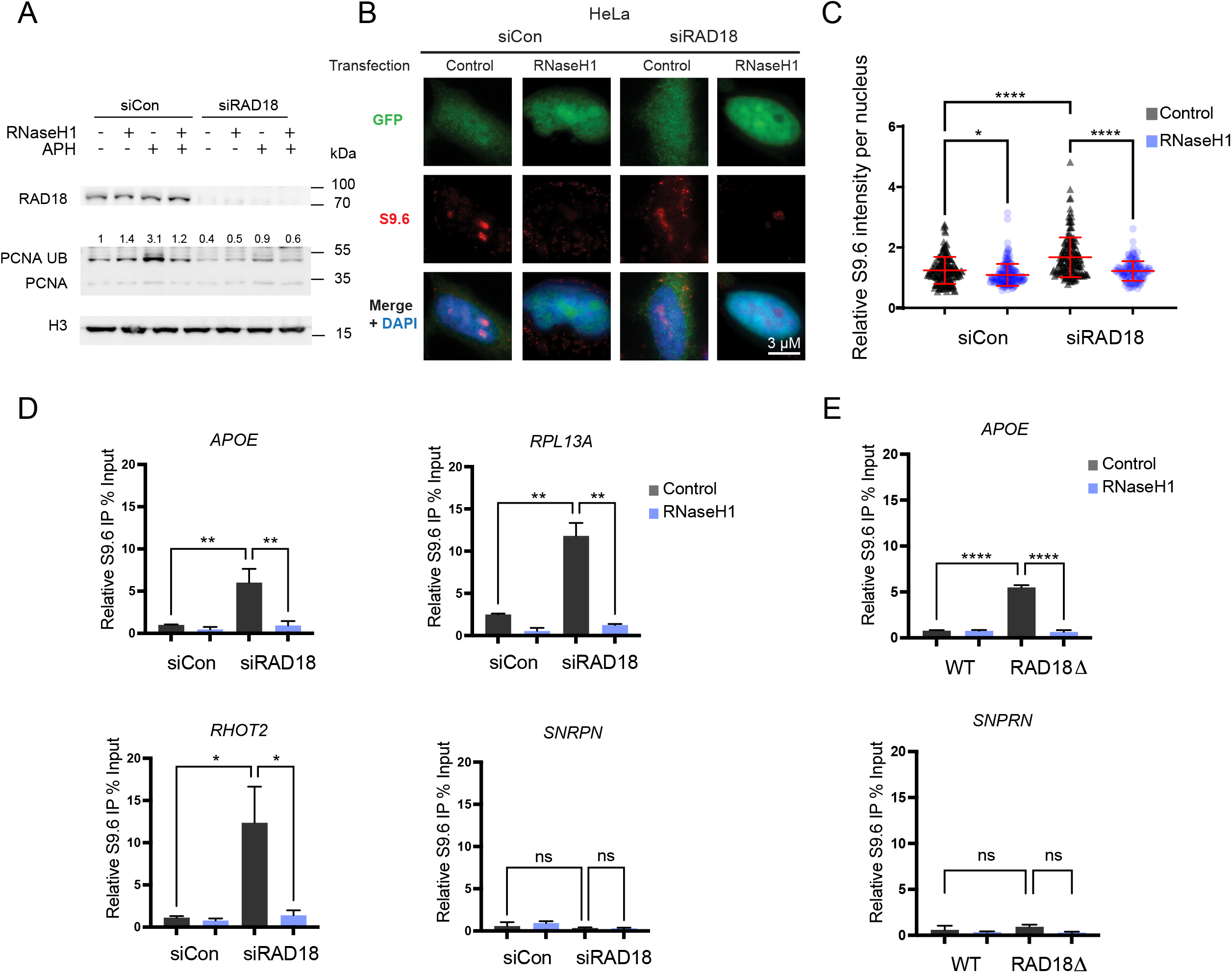
Loss of RAD18 increases DNA:RNA hybrids. (A) Western blot of chromatin bound lysates representing RAD18 knockdown with pooled siRNA and the subsequent reduction of PCNA ubiquitination in HeLa cells. Ubiquitinated PCNA is increased in cells treated with APH for 24 hours, and this is reduced in cells transfected with RnaseH1 expression plasmid. (B,C) Nuclear S9.6 antibody staining in non-targeting control siRNA or siRAD18 treated HeLa cells with or without transfection of GFP-tagged RNaseH1 overexpression plasmid. B, representative images, C, quantification. (D) DRIP-qPCR in non-targeting control siRNA and siRAD18 HeLa cells at R-loop prone sites APOE, RPL13A, RHOT2, and negative control site SNPRN with and without RNaseH1 treatment. (E) DRIP-qPCR in P53^-/-^ RPE1-hTERT WT and RAD18 knockout clone cells at R-loop prone sites APOE and negative control site SNPRN. Treatment concentrations of APH (400 nM). ANOVA p<0.05, N=3 for all. Error bars represent standard deviation. *siCon*, non-targeting control siRNA; *ns*, not significant; *, p<0.05; **, p<0.01; ***, p<0.001; ****, p<0.0001.

### Yeast growth and recombination assay

Trp1- derivatives of BY4741 yeast or a *rad18Δ::KanMX* derivative were grown in standard synthetic complete media at 30°C. These strains carried an empty pRS316-URA3 control plasmid or one expressing yeast RNH1 from its native promoter. Recombination frequencies in the direct-repeat plasmids pARSHLB-IN, pARSHLB-OUT (gifts of A. Aguilera, CABIMER, Sevilla, Spain), were determined by the number of LEU+ colonies emerging in the total population of TRP+ plasmid transformants as described (31, 32).

### Immunofluorescence

For all immunofluorescence experiments, cells were grown on coverslips overnight before siRNA transfection and plasmid overexpression. For S9.6 immunofluorescence, cells were washed with PBS, fixed with ice-cold methanol for 10 min and permeabilized with ice-cold acetone for 1 min. After PBS wash, cells were blocked in 3% bovine serum albumin (BSA), 0.1% Tween-20 in 4× saline sodium citrate buffer (SSC) for 1 h at room temperature. Cells were then incubated with primary antibody S9.6 (1:500, S9.6, Kerafast) and in certain experiments nucleolin (1:1000, Abcam) overnight at 4 °C. Cells were then washed three times in PBS and stained with mouse (rabbit for nucleolin) Alexa-Fluor-488 or 568-conjugated secondary antibody (1:1000, Life Technologies) for 1 h at room temperature, washed three times in PBS, and stained with DAPI for 5 min. Cells were imaged on a LeicaDMI8 microscope at ×100 and ImageJ was used for processing and quantifying nuclear S9.6 intensity in images. For experiments with GFP overexpression, only GFP-positive cells were quantified. For phospho-ATM (1:1000, ser1981, (D25E5) 13050, cell signaling), γH2AX (1:1000, H2A.X (phospho-S139) [EP854(2)Y], ab81299, abcam), RAD18 (1:1000, NB100-61063, Novus) and FANCD2 (1:1000, NB100-182SS, Novus) the immunostaining was performed instead with 4% paraformaldehyde for 15 min and permeabilization with 0.2% Triton X-100 for 5 min. For EU (ethynyl uridine) staining, cells grew grown on coverslps as mentioned above, treated with triptolide (40 nM) for 2 hours, and the Click-iT® RNA Imaging Assay (Thermo Fisher) was used according to manufacturer instructions.

### Western blotting

Whole-cell lysates were prepared with RIPA buffer containing protease inhibitor (Sigma) and phosphatase inhibitor (Roche Applied Science) cocktail tablets and the protein concentration were determined by Bio Rad Protein assay (Bio-Rad). Equivalent amounts of protein were resolved by SDS-PAGE and transferred to polyvinylidene fluoride microporous membrane (Millipore), blocked with 1.5% BSA in H20 containing 0.1% Tween-20 (TBS-T), and membranes were probed with the following primary antibodies: anti-RAD18 (Bethyl A301-340A) (1:1000), anti-PCNA (Abcam [PC10] ab29) (1:1000), anti-ubiquityl-PCNA (Lys164) (NE Biolabs 13439S) (1:1000), anti-GAPDH (Thermofisher MA5-15738) (1:2000), anti-FANCD2 (Novus, NB100-316), anti-Vinculin (Santa Cruz, sc-25336), anti-H3 (Cell Signaling, 9715). Secondary antibodies were conjugated to horseradish peroxidase (HRP) and peroxidase activity was visualized using Chemiluminescent HRP substrate (Thermo Scientific). Western blot densitometry analysis was performed using ImageJ and are expressed relative to the first lane control and normalized to the indicated loading control. For chromatin fractionation, cells were washed twice with ice-cold PBS and prepared with ice-cold 1% TritonX-100 in PBS. Lysates were vortexed, incubated on ice for 15 minutes, and centrifuged at 4°C. Pellet was resuspended in 0.2N HCl, vortexed, incubated on ice for 20 minutes and centrifuged at 4°C. Supernatant was neutralized with the 1 M Tris-HCL pH 8.

### Monitoring plasmid copy number in transcription-replication conflict system

HEK293 cells expressing the tetracycline (Tet)-regulated transactivator were transfected using Lipofectamine 3000 (Invitrogen) with doxycycline (DOX) (Sigma) inducible plasmids containing one of four plasmids containing either mAIRN or ECFP genes with promoters oriented head-on (HO) or co-directional (CD) with respect to replication; ECFP-HO, ECFP-CD, mAIRN-HO, and mAIRN-CD (**Table S2**). Once cells were stable expressing plasmids, plasmid copy number was measured with PCR using primers specific for the plasmid or for using an actin control to measure cell count (**Table S1**). Upon transcription induction with 1000 ng/mL DOX for 72 hours, cells were 2-4 × 10^5 cells were lysed and DNA was extracted using the same method as the Hamperl *et al*. qPCR was performed on AB Step One Plus real-time PCR machine (Applied Biosystems) using Fast SYBR Green Master Mix (Applied Biosystems) and plasmid copy number was represented as the ratio of the amount of oriP to the amount of beta-Actin.

### Comet Assay

The neutral comet assay was performed using the CometAssay Reagent Kit for Single Cell Gel Electrophoresis Assay (Trevigen) in accordance with the manufacturer’s instructions. In brief, cells were combined with LMAgarose at 37 °C at a ratio of 1:10 (cell:agarose) and spread onto CometSlide. After gelling at 4°C in the dark, the slides were then immersed in 4°C Lysis Solution overnight. The next day, slides were removed from Lysis Solution and immersed in 4°C 1 × Neutral Electrophoresis Buffer for 30 min. Electrophoresis was then performed at 4°C at 20 Volts for 30 min. Slides were then immersed in DNA Precipitation Solution for 30 min followed by 70% ethanol for 30 min at room temperature, and dried at 37°C for 10 min. Finally, slides were stained with PI and imaged on LeicaDMI8 microscope at ×20. Comet tail moments were obtained using an ImageJ plugin as previously described (33). At least 50 cells per sample were analyzed from each independent experiment.

### Proximity Ligation Assay

The proximity ligation assay was performed using the Duolink In Situ Kit (Sigma) in accordance with the manufacturer’s instructions. Cells were seeded on 18 mm circular slides, washed with PBS twice, fixed with 4% PFA in PBS for 10 min and permeabilized with 0.2% Triton X-100 for 10 min. Cells were then blocked in 3% BSA, 0.1% Tween-20 in 4X SSC for 1 h at room temperature. Cells were then incubated with primary antibody overnight at 4 °C [1:500 goat anti-RNA polymerase II antibody (PLA0292, Sigma) with 1:500 rabbit anti-PCNA antibody (PLA0079, Sigma)]. The next day cells were washed 2x in PBS and then incubated with pre-mixed PLA probe anti-goat minus and PLA probe anti-rabbit plus (Sigma) for 1 h at 37 °C. The following ligation and amplification detections steps were followed identically to the Duolink In Situ Kit (sigma). Slides were then stained with DAPI and imaged on a LeicaDM18 microscope at ×100. Negative controls were treated identically but anti-RNA polymerase II antibody and PCNA antibody were omitted.

### EdU Click-IT Imaging

EdU incorporation into cells was performed using the Click-IT™ EdU Cell Proliferation Kit for Imaging, Alexa Fluor™ 647 (Thermofisher) in accordance with the manufacturer’s instructions. Immunofluoresence was performed as listed above.

### SIRF: in situ protein interactions at nascent and stalled replication forks

SIRF was carried out as previously described (34). Cells were seeded on slides and incubated with 125 μM EdU for 8 min except for the negative control. After EdU was removed and cells were washed with PBS for two times, fresh growth media was replaced with the specific drug containing media. After 2 hour drug treatment, cells were washed with PBS two times before fixation with 3% PFA in PBS and permeabilized with 0.25% TritonX-100 in PBS. After blocking, cells were incubated with the following primary antibodies overnight: rabbit anti-RAD18 NB100-61063 (Novus) (1:200) and mouse anti-biotin (D5A7) (1:200) (Cell signaling). The rest of the protocol follows Proximity Ligation Assay as above.

### DRIP and ChIP qPCR

ChIP was performed using the SimpleCHIP enzymatic chromatin IP kit (Cell Signaling Technologies) in accordance with manufacturer’s instructions with some modifications. HeLa cells were seeded at 4×10^6^ in 15 cm plates and treated with either DMSO or 400 nM APH for 24 hours. Cells were crosslinked in 1% formaldehyde for 10 minutes, followed by 2 minute incubation in 1 x glycine solution (CST#7005). Cells were washed in 20 mL cold PBS on ice twice and scrape cells down into 1 ml ice-cold 1X Buffer A (in 500 µM DTT and 1 x protease inhibitor cocktail (PIC) (CST#7012) per IP prep in sonication tubes. Nuclei were pelleted and resuspended in 300 ul of 1 x ChIP buffer and 1 x PIC and incubated on ice for 10 minutes. DNA were sonicated on Q Sonica Sonicator Q700 for 8 min (20 sec ON, 30 s OFF) at 100 µm amplitude to fragment DNA. Lysates were clarified with centrifugation and the supernatant was collected and split into ChIP analysis, DRIP and DNA content measurements. ChIP samples were stored at -80°C until future use. For DRIP, DNA samples were treated with RNAse A (CST#7013) followed by proteinase K (CST#10012) overnight at 65°C. DNA samples were purified with Cell Signaling spin columns (#14209S). DNA concentration was measured using NanoDrop One/Onec. DNA samples were used for DRIP analysis, where 2% input sample was isolated and DNA was separated into no treatment and RNaseH treatment. 1 μl of 5000 units/ml RNaseH (NEB, M0297L) was used per µg of DNA and incubated for 48 hours at 37°C. ChIP-Grade protein G magnetic beads (Cell Signaling, 9006S), 25 μl per IP, were preblocked in preblocking buffer (PBS/EDTA containing 0.5% BSA) for 1 hour rotating at 4°C. Beads were immobilized with S9.6 antibody (MABE1095) using 5 µg per IP, in 1 x ChIP buffer and 1 x PIC for 4 hours rotating at 4°C followed by addition of DNA and incubated overnight rotating at 4°C. Samples were washed 3 x low salt buffer, 1 x with high salt buffer, and eluted with 1 x elution buffer (CST#7009) for 30 minutes at 65°C vortexing intermittently. Beads were pelleted and supernatant was purified with Cell Signaling spin columns (CST#14209S). qPCR primer sequences can be found in **Table S1**. qPCR was performed on AB Step One Plus real-time PCR machine (Applied Biosystems) using Fast SYBR Green Master Mix (Applied Biosystems). ChIP analysis for FANCD2 was performed similarly, however, antibodies are added directly to digested crosslinked chromatin and incubated overnight rotating at 4°C. Preblocked beads are then added to chromatin-antibody samples for 4 hours rotating at 4°C followed by the same washes and elution procedures. Samples were then treated with RNAse A (CST#7013) followed by proteinase K (CST#10012) overnight at 65°C, purified with Cell Signaling spin columns (CST#14209S), and qPCR was performed on AB Step One Plus real-time PCR machine (Applied Biosystems) using Fast SYBR Green Master Mix (Applied Biosystems).

## RESULTS

### Loss of RAD18 increases DNA:RNA hybrids

Our previous work identified yeast Rad18 as a potential regulator of R-loops in a screen of yeast DNA repair mutants with increased staining for the S9.6 DNA:RNA hybrid antibody (30). A similar screen performed with the S9.6 antibody in HeLa cells also identified RAD18 as a candidate hybrid regulator (35). Based on these data, we set out to test a role for RAD18 in tolerating DNA:RNA hybrids in a human cell model. RAD18 protein was depleted in HeLa cells by using a pool of four siRNAs to limit off-target effects. To validate our RAD18 knockdown system, we tested loss of PCNA ubiquitination activity in RAD18 deficient HeLa cells. Western blot analysis showed RAD18 deficient cells had lower chromatin bound ubiquitinated PCNA using a ubiquitin-specific PCNA antibody (**Figure 1A**). We transfected cells with GFP-tagged RNaseH1, an endoribonuclease that hydrolyzes the phosphodiester bond of RNA hybridized to DNA, to effectively degrade R-loops in this system. RNaseH1 did not decrease PCNA ubiquitination in unstressed cells, therefore we treated cells with APH, a DNA polymerase inhibitor. Low doses of APH blocks a subset of replication, promoting conflicts and R-loop formation, especially at long genes in common fragile sites (CFS) (36). APH treated cells increased PCNA ubiquitination specifically dependent on RAD18 status (**Figure 1A**). Furthermore APH-induced PCNA ubiquitination was reduced in RNaseH1 treated HeLa cells, suggesting R-loops are responsible for triggering this activation. DNA:RNA hybrid levels were measured with S9.6 immunofluorescence combined with RNaseH overexpression controls (**Figure 1B** and quantified in **Figure 1C**). An increased level of RNaseH-sensitive S9.6 signal was detected in RAD18 deficient cells with no external stress, suggesting that R-loops are stabilized when RAD18 is absent. We confirmed this finding in CRISPR-generated RAD18 knockout clones of P53^-/-^ RPE1-hTERT cells (**Figure S1A, S1C-E**). Given the cross-reactivity of S9.6 for ribosomal RNA in imaging experiments (37), we also tested DNA:RNA immunoprecipitation (DRIP) in RAD18 depleted cells. We found a significant RNaseH-sensitive increase in DRIP signal at APOE, RPL13A and RHOT2, each previously characterized loci containing DNA sequences prone to R-loop formation (35) **(Figure 1D**). Analysis of DRIP-qPCR accumulation in knockout clones confirmed an increase in signal when RAD18 was absent (**Figure 1E**). Together these data show that, as suggested in other systems, RAD18 depletion or knockout increases R-loop accumulation.

### RAD18 localization is dependent on R-loop formation and transcriptional activity

RAD18 is a central regulator of PCNA through its localization to sites of replication stress where it initiates signaling by PCNA ubiquitination (6). We found that RAD18 foci formed spontaneously in HeLa cells and that RNaseH1 overexpression significantly reduced RAD18 foci levels (**Figure 2A** and quantified in **Figure 2B**). Furthermore, treatment of cells with 40 nM of triptolide for 2 hours, which reduces global transcription (**Figure S1B**), reduced RAD18 foci formation (**Figure S2A**). Alternatively, a 24 hour treatment of low-dose APH further increased RAD18 foci formation (**Figure S2A)**. This suggests that R-loops and transcription may constitute an ongoing stress that recruits RAD18 to foci in these cells. RAD18 foci presumably reflect recruitment to stressed replication forks, and we utilized the SIRF assay (Quantitative in situ analysis of protein interactions at DNA replication forks) (38), to confirm that RAD18 localizes to replicating DNA based on proximity-ligation assay (PLA) between EDU incorporated into DNA and RAD18 (**Figure S2B**). To determine whether there was an association between RAD18 and R-loop prone sites, we precipitated RAD18 for ChIP-qPCR analysis at the previously tested APOE, RPL13A and RHOT2 sites. This showed that RAD18 pulldown is strongly enriched at R-loop prone sites. Additionally, triptolide treatment significantly reduced RAD18 binding to these sites but not a control site SNPRN not susceptible for R-loop formation (**Figure 2C**). Recruitment of RAD18 to R-loops was further interrogated using a PLA experiment between RAD18 and S9.6 with APH and triptolide treatment in the P53^-/-^ RPE1 cells (**Figure S2C**). Similar to RAD18 foci, RAD18-S9.6 PLA showed that APH treatment increased foci formation, however triptolide treatment did not significantly decrease PLA foci. One reason for this could be that the endogenous stress caused by R-loops and conflicts is expected to be lower in an RPE1 background, especially in cells with functioning RAD18, as these cells are relatively stable. RAD18 KO cells acted as a negative control with near zero foci (**Figure S2C**). These data suggest that RAD18 accumulates in foci and at specific genomic loci in a manner dependent on R-loops and transcription, respectively.

**Figure 2.**
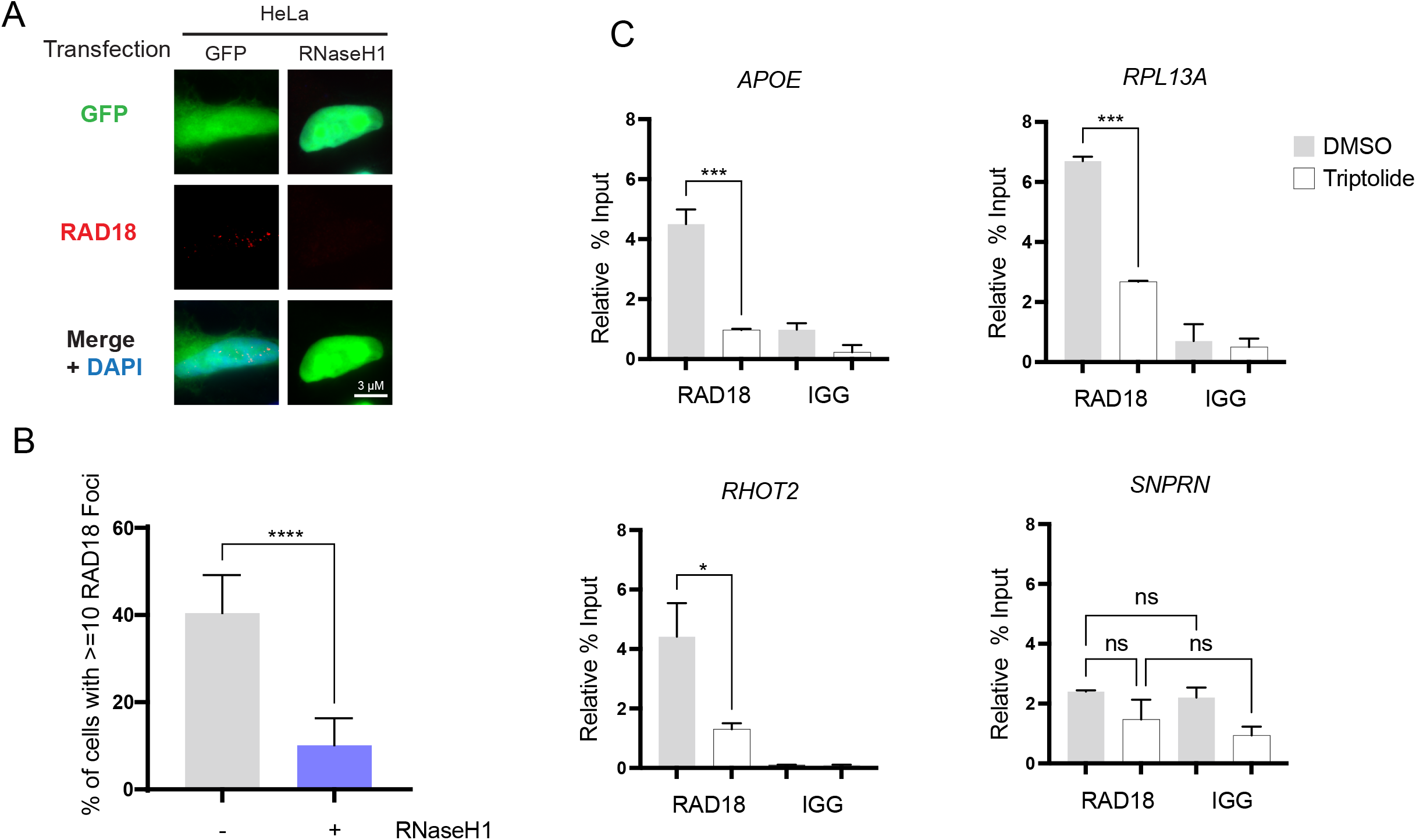
RAD18 localization is dependent on R-loop formation and transcriptional activity. (A-B) Suppression of spontaneous RAD18 foci in GFP-tagged RNaseH1 transfected cells. A, representative images, B, quantification. (C) RAD18 ChIP in Hela cells at R-loop prone sites APOE, RPL13A, RHOT2, and negative control site SNPRN with and without a 2 hour treatment of triptolide and compared to an IGG negative control. Treatment concentrations of triptolide (40 nM). (B) Student T Test (C) ANOVA. p<0.05, N=3 for all. *ns*, not significant; *, p<0.05; **, p<0.01; ***, p<0.001; ****, p<0.0001.

### Loss of RAD18 increases DNA damaging transcription-replication conflicts

While RAD18 is a replication fork associated protein (**Figure S2B**), our data so far suggest a role for transcription in its recruitment to R-loop sites. This suggests that transcription-replication conflicts are a likely contributor to the observed R-loop accumulation. To test this we first took advantage of the published PLA technique monitoring PCNA and RNA polymerase II proximity in cells (18). Depletion of RAD18 strongly induced PCNA-RNAPII PLA foci, and this increase was reversible when RNaseH1 was overexpressed (**Figure 3A** and quantified in **3B**). This finding was surprising in light of previous work suggesting that ssDNA gaps left in RAD18-depleted cells may be substrates for R-loop formation after the replisome has passed (35). This model would suggest that no conflicts, or co-directional conflicts would arise in RAD18-depleted cells. Since our PLA assay does not tell us about the direction of conflicts, we used two more assays to determine whether head-on (HO) or co-directional (CD) collisions were more damaging. First we used a set of direct repeat recombination reporter plasmids in yeast with opposite transcript orientation. This assay showed that deletion of yeast RAD18 enhanced recombination in both CD and HO orientation, but that the magnitude was great in HO conflicts (**Figure 3C**). Co-expression of RNaseH1 reduced conflicts to WT levels in both cases, supporting a role for R-loops in RAD18 mutants across species (**Figure 3C**). Next we used a human conflict plasmid system developed by the Cimprich lab to test the effects in RAD18-depleted HeLa cells (18). In this system an R-loop prone (mAIRN) or control (ECFP) sequence is transcribed in the head-on (HO) or co directional (CD) orientation with respect to replication, and plasmid copy number is monitored by qPCR (**Table S2**). Plasmid instability is represented by decreased copy number in this assay and represents a higher degree of plasmids with unresolved conflicts. We performed this experiment in control or RAD18-depleted cells and found a significant reduction in stability in the HO and R-loop prone sequence plasmid when RAD18 was absent (**Figure 3D-E**). Overall, our data suggest that transcription-replication conflicts accumulate in cells lacking RAD18, highlighting the role of transcription as an endogenous source of stress.

**Figure 3.**
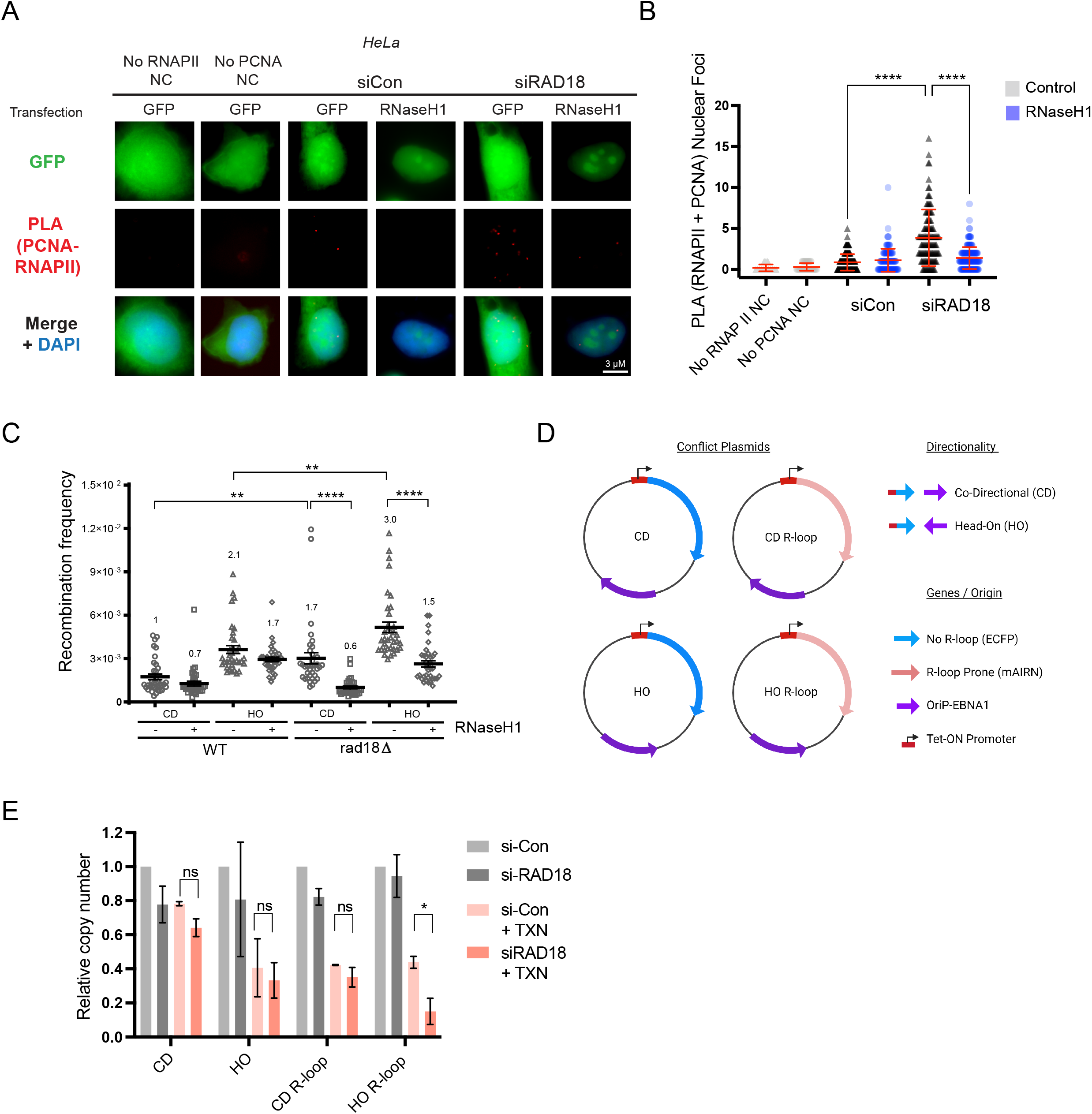
Loss of RAD18 increases DNA damaging transcription-replication conflicts. (A-B) Transcription-replication conflicts measured by RNAPII-PCNA proximity ligation in non-targeting control siRNA or siRAD18 treated HeLa cells with or without transfection of a GFP-tagged RNaseH1 overexpression plasmid. A, representative images, B, quantification. (C) *LEU2* direct repeat recombination frequencies in the indicated yeast strains with (+) or without (-) ectopic RNaseH1 expression. Plasmids transcribe the reporter in S-phase in either a head-on (HO) or codirectional (CD) with respect to replication. (D,E) Plasmid-based head-on or co-directional conflict stability assay, a measure of plasmid copy number with or without doxycycline-induced transcription activation on ECFP (R-loop negative) and mAIRN (R-loop prone) constructs in non-targeting control siRNA or siRAD18 treated HeLa cells. D, schematic. E, quantification. ANOVA p<0.05, N=3 for all. Error bars represent standard deviation. *PLA*, proximity ligation assay; *siCon*, non-targeting control siRNA; *HO*, head-on conflict; *CD*, codirectional conflict; *TXN*, 72 hour transcription induced with 1 µg/ml doxycycline; *No RNAPII NC*, negative control lacking the RNA Polymerase II antibody; *No PCNA NC*, negative control lacking the PCNA antibody; *ns*, not significant; *, p<0.05; **, p<0.01; ***, p<0.001; ****, p<0.0001.

### RAD18 suppresses transcription- and R-loop-dependent DNA replication stress

Elevated conflicts are associated with DNA replication stress (39) and we therefore predicted that RAD18 depletion would drive increased R-loop associated replication stress. To test this we stained siRAD18 or control cells with an antibody for RPA2-ser33-phosphorylation which is deposited on RPA during replication stress (40). As predicted, RAD18 depleted cells had an elevated number of RPA2-Ser33P foci, even with no external stressors. Ectopic expression of RNaseH1 decreased RPA2-Ser33P foci levels in both control and siRAD18 cells (**Figure 4A** and quantified in **Figure 4B**). R-loops causing replication stress, or stalled replication forks themselves, can be processed by nucleases to give rise to DNA double strand breaks (DSBs) (41,42). To assess the impact of R-loops on DNA damage in our RAD18 knockdown model, we used immunofluorescence of phosphorylated H2AX (γH2AX) to detect DNA DSBs. As expected, RAD18 depletion led to a greater number of cells with γH2AX foci (**Figure 4C** and quantified in **Figure 4D**). Overexpression of RNaseH1 returned the percentage of cells with γH2AX foci back to control levels, implicating R-loop accumulation as the source of DNA damage in RAD18 deficient cells. This same trend of both enhanced RNaseH-sensitive replication stress and DNA damage was further confirmed in the RAD18 KO cell line (**Figure S3A-B)**. In parallel, neutral comet assays were performed to directly detect DNA DSBs in RAD18 depleted cells. Again, DSB increases in siRAD18 cells were partly rescued by RNaseH1 overexpression (**Figure S3C** and quantified in **Figure S3D**). To confirm the importance of global transcription as a driver of replication stress and DNA damage in RAD18-deficient cells, we used the RPE1 knockout model for immunofluorescence experiments of phosphorylated-ATM and RPA2-ser33-phosphorylation. These data show the accumulation of spontaneous damage and replication stress in RAD18-KO cells, marked by phospho-ATM and RPA2-Ser33P respectively, is significantly reduced by pre-treatment with triptolide (**Figure S3E** and quantified in **Figure S3F-G**). To implicate replication as a source of this DNA damage, we used an EdU click-IT kit to identify which cells are undergoing DNA synthesis. Interestingly, we observed a higher proportion of RAD18 knockout cells in S-phase, suggesting that endogenous DNA damage could be triggering the intra-S-phase checkpoint (**Figure 4E**). In the P53^-/-^ RPE1-hTERT RAD18 KO cells, we identified that γH2AX foci formation is predominantly occurring in EdU positive cells (**Figure 4F)**. RAD18 has multiple roles in tolerating genotoxic stress, most of which require RAD18 E3 ubiquitin ligase activity (5). To establish if RAD18-dependent PCNA ubiquitination is important for R-loop accumulation, we used PCNA^K164R^ MEFs expressing a non-ubiquitylable PCNA. We found that the mutant had higher S9.6 signal compared to the parental cell line. Importantly, as seen in human cell lines, RAD18 knockdown increased S9.6 in parental MEFs, however RAD19 depletion did not result in any further increases in staining in the PCNA^K164R^ mutant (**Figure S4A**). This suggests that the ubiquitination of PCNA is in the same pathway as RAD18. Together, these results support the hypothesis that accumulated R-loops and conflicts in RAD18 deficient cells are drivers of DNA damage and replication stress and that RAD18 E3 ubiquitin ligase activity is likely required.

**Figure 4.**
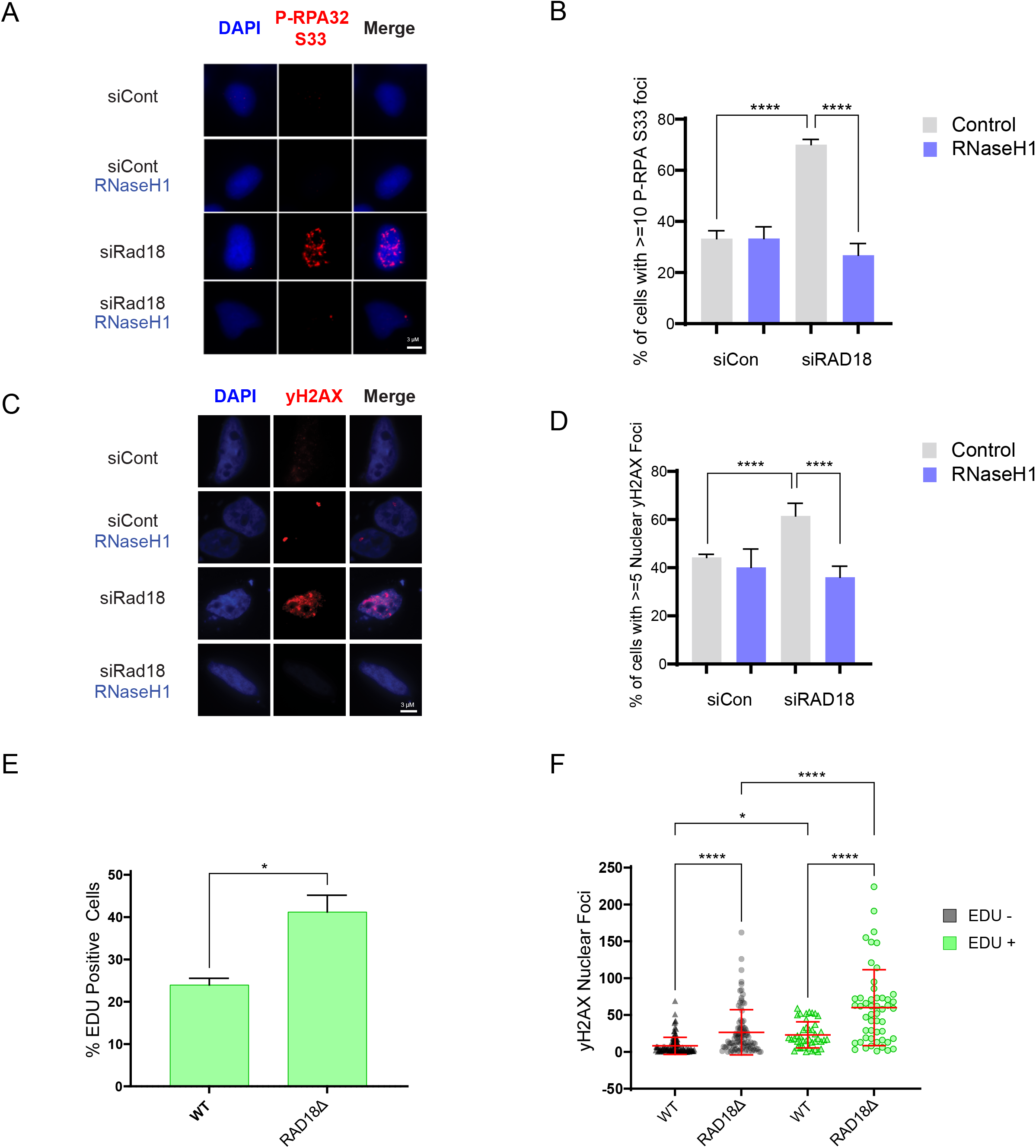
RAD18 suppresses transcription and R-loop associated DNA replication stress and DNA damage. (A,B) RPA2-ser33 phosphorylation immunofluorescence in non-targeting control or siRAD18 treated HeLa cells with or without transfection of a GFP-tagged RNaseH1 overexpression plasmid. A, representative images, B, quantification. (C,D) H2AX-ser139 phosphorylation immunofluorescence in non-targeting control or siRAD18 treated HeLa cells with or without transfection of RNaseH1 overexpression plasmid. C, representative image, D, quantification. (E) Percentage of cells exhibiting EdU fluorescence in P53^-/-^ RPE1-hTERT WT and RAD18 knockout clone cells. (F) H2AX-ser139 phosphorylation immunofluorescence in P53^-/-^ RPE1-hTERT WT and RAD18 knockout clone cells in both EdU negative and positive cells. Treatment concentrations of triptolide (40 nM) and EdU (10 µM). (B,D) Fisher’s Exact Test, p<0.05, n=3. (E) Student T Test, p<0.05, n=3. (F) ANOVA, p<0.05, n=3. Error bars represent standard deviation for all. *APH*, aphidicolin; *PladB*, pladienolide B; *EDU*, Ethynyl-2’-deoxyuridine; *siCon*, non-targeting control siRNA; *ns*, not significant; *, p<0.05; **, p<0.01; ***, p<0.001; ****, p<0.0001.

### RAD18-dependent FANCD2 foci recruitment to R-loop stressed sites

RAD18-dependent PCNA ubiquitination coordinates TLS, fork reversal, template switching, and potentially FA pathway activation. Many reports have suggested that the FA pathway is a key player involved in TRC and R-loop resolution and tolerance (24,30,43). In contrast, direct tests of TLS polymerase knockdown failed to show increased R-loops in previous studies, although RNaseH1 dependent DNA damage was observed (44). Therefore, we hypothesized that effects on the FA pathway could explain the RAD18 knockdown phenotypes, despite FA activation by RAD18-PCNA being less well understood. To test the hypothesis that RAD18 is important for FANCD2-dependent R-loop and conflict regulation, we wanted to distinguish between the role of the FA pathway in tolerating DNA:RNA hybrids and its role in resolving ICLs. To do this we used the crosslinker MMC to induce ICLs, or the U2 spliceosome inhibitor Pladienolide B (PladB) to induce R-loops. Disruption of SF3B1 in the U2 spliceosome results in a redistribution of R-loops throughout the genome through increased promoter pausing and decreased elongation, potentially resulting in increased R-loops at readthrough transcription events (https://www.biorxiv.org/content/10.1101/2020.06.08.130583v1). Interestingly, either treatment with the crosslinker MMC or with PladB resulted in an increase in PCNA-RNAPII PLA foci, supporting the view that different types of stress lead to conflicts (**Figure 5A**), consistent with our observations in Figure 4. As expected, we also found that cells deficient in FANCD2 had higher PCNA-RNAPII PLA foci that were restored to normal levels in cells overexpressing RNaseH1 (**Figure S4B**). Next we tested FANCD2 colocalization with R-loops using PLA. FANCD2-S9.6 PLA signal was evident in unstressed cells, reiterating that FANCD2 colocalizes with DNA:RNA structures (**Figure 5B**). Importantly, MMC treatment had no effect on FANCD2-S9.6 PLA, while PladB treatment significantly increased the signal (**Figure 5B**). This finding is reinforced by RAD18-S9.6 PLA levels that increased in PladB treated cells but not MMC treated cells (**Figure S2D**). In HeLa cells, FANCD2 foci formation was stimulated both by PladB and MMC treatment, highlighting the dual role of FANCD2 and the FA pathway in tolerating both R-loops and ICLs (**Figure 5C**). To analyze the importance of RAD18 in coordinating FANCD2 in the presence of both R-loops and ICLs we depleted RAD18 or the FANCD2-targeting E3 ubiquitin ligase FANCL using siRNA. FANCL knockdown significantly reduced FANCD2 foci formation under both conditions (**Figure 5C**). RAD18 depletion was as penetrant as FANCL in reducing PladB induced FANCD2 foci, but had significantly weaker reductions in FANCD2 foci after MMC treatment (**Figure 5C**). This suggests that RAD18 has a critical role in FANCD2 recruitment during R-loop associated replication stress, but a partially dispensable role, compared with FANCL, when responding to ICLs. To test the effects of RAD18 on a better characterized stressor linked to R-loop TRCs, we tested the effects of low dose APH on FANCD2 foci. A 24 hour treatment of low dose APH strongly induced FANCD2 foci formation in HeLa cells in a manner dependent on RAD18 and FANCL (**Figure 5D**). To confirm these findings, FANCD2 foci were measured in the RAD18 knockout clones of P53^-/-^ RPE1-hTERT cells using PladB and APH treatment (**Figure 5E** and quantified in **Figure 5F**). Similar to the RAD18 and FANCL knockdown findings, we found a smaller percentage of FANCD2 foci positive cells in RAD18 knockout cells (**Figure 5F**). Furthermore, we found that treatment with these stressors resulted in a higher degree of DNA damage, represented by phospho-ATM foci, in the RAD18 knockout cells (**Figure S5A** and quantified in **Figure S5B**). Additionally, FANCD2-S9.6 PLA levels in P53^-/-^ RPE1-hTERT WT cells were reduced in triptolide and MMC treated cells and, APH treatment increased FANCD2-S9.6 PLA specifically in WT cells and not RAD18 KO cells, implicating the requirement of RAD18 for FANCD2 R-loop localization **(Figure S5C)**. Overall these data suggest a more important role for RAD18 in FANCD2 recruitment to transcription-associated stresses than to ICLs. RAD18 ubiquitination requires binding of the E2 ubiquitin-conjugating protein RAD6 for its catalytic activity. To test if the E3 ubiquitin ligase activity of RAD18 was required for the APH induced foci formation of FANCD2, we transfected the RAD18 KO clones with either full-length RAD18 plasmid, or with RAD6-binding domain mutant plasmid (**Table S2**)(**Figure S5D**) (45). We found that transfection of full-length RAD18, but not of the RAD6-binding mutant, rescued the APH induced foci formation, implicating the RAD18 E3 ligase activity in FANCD2 recruitment during replication stress (**Figure 5G** and quantified in **Figure 5H)**. This data reiterates that FANCD2 activation is induced in multiple contexts of cellular stress, highlighted by either ICL or R-loop induction. Furthermore, RAD18 is important for coordinating both pathways, perhaps more strongly contributing to recruitment in response to R-loops and conflicts through its E3-ubiquitin ligase activity.

**Figure 5.**
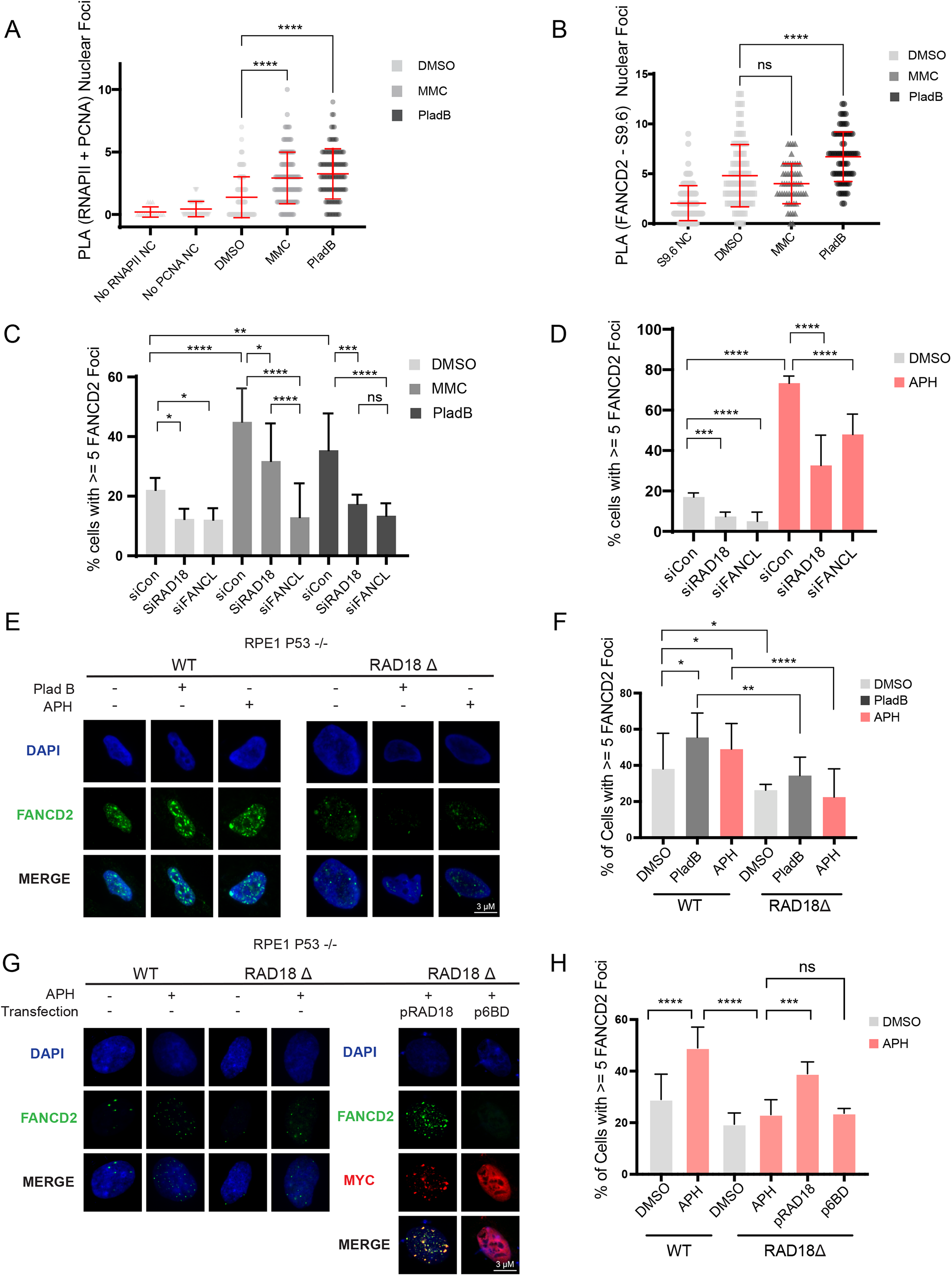
RAD18 mediates recruitment of FANCD2 to a subset of replicative lesions. (A) Proximity of RNAP II and PCNA in HeLa cells treated with DMSO, Pladienolide B or Mitomycin C for 2 hours using the proximity ligation assay. (B) Proximity of FANCD2 and S9.6 in HeLa cells treated with DMSO, Pladienolide B or Mitomycin C for 2 hours using the proximity ligation assay. (C) FANCD2 immunofluorescence of non-targeting control siRNA, siRAD18, or siFANCL HeLa cells treated with DMSO, Pladienolide B or Mitomycin C for 2 hours. (D) FANCD2 immunofluorescence of non-targeting control siRNA, siRAD18, or siFANCL HeLa cells treated with DMSO or APH for 24 hours. (E,F) FANCD2 foci immunofluorescence in P53^-/-^ RPE1-hTERT RAD18-KO and WT cells treated with either DMSO, Pladienolide B for 2 hours or APH for 24 hours. E representative image, F quantification. (G,H) FANCD2 foci imaging in P53^-/-^ RPE1-hTERT RAD18-KO cells rescued with WT (pRAD18) or Rad6-binding mutant (p6BD) plasmids compared to WT. Cells transfected with MYC-tagged rescue constructs were identified and scored by anti-MYC staining as shown in G. G representative images, H quantification. Treatment concentrations of Pladienolide B (5 µM), Mitomycin C (1.5 µM) and APH (400 nM). (A-B) ANOVA, p<0.05, n=3. (C,D,F,H) Fisher’s Exact Test, p<0.05, n=3. Error bars represent standard deviation for all. *PLA*, proximity ligation assay; *siCon*, non-targeting control siRNA; *MMC*, mitomycin C, *PladB*, pladienolide B, *APH*, aphidicolin; *No RNAPII NC*, negative control lacking the RNA Polymerase II antibody; *No PCNA NC*, negative control lacking the PCNA antibody; *ns*, not significant; *ns*, not significant; *, p<0.05; **, p<0.01; ***, p<0.001; ****, p<0.0001.

### FANCD2 Recruitment to CFS and R-loop sites is Dependent on RAD18

Finally, to evaluate the impact of R-loop accumulation and FANCD2 recruitment at specific genomic loci prone to transcription-replication conflicts, we first investigated CFSs (46). Genes within CFSs are typically much larger than average and may take more time than a single cell cycle to transcribe, resulting in transcription-replication conflicts and R-loop stabilization (47). Additionally, CFSs become unstable in the presence of replication stress. Treatment with low concentrations of APH is known to result in FANCD2 accumulation at R-loops and CFS regions (48). FANCD2 has been specifically linked as a trans-activating facilitator of CFS replication, reducing the number of R-loops at CFSs and promoting unperturbed replication (49). We used DRIP-qPCR to monitor R-loop accumulation after a 24-hour APH treatment. This showed a significantly higher DRIP signal in APH treated cells, which was further increased in RAD18 deficient cells at the NRG3 CFS (**Figure 6A**). Indeed, RAD18 ChIP to the fragile site at NRG3 was strongly induced by APH treatment (**Figure 6B**), consistent with the accumulation of stalled replication forks in this region. Since RAD18 depletion strongly reduced FANCD2 foci after APH treatment (**Figure 5**), we next assessed FANCD2 binding to CFSs at NRG3, WWOX and CDH13 loci after APH treatment. We found that FANCD2 strongly accumulates at each CFS loci after APH treatment in control siRNA treated HeLa cells but that cells depleted for RAD18, FANCD2 accumulation was significantly reduced (**Figure 6C** and **Figure S6A-B)**. Additionally, FANCD2 binding to R-loop prone sites at APOE and RPL13A was also dependent on RAD18 (**Figure S6C-D)**. These data support a model in which RAD18 initiates a cascade of events leading to recruitment of FANCD2 at R-loop prone and transcription-associated fragile sites (**Figure 6D**).

**Figure 6.**
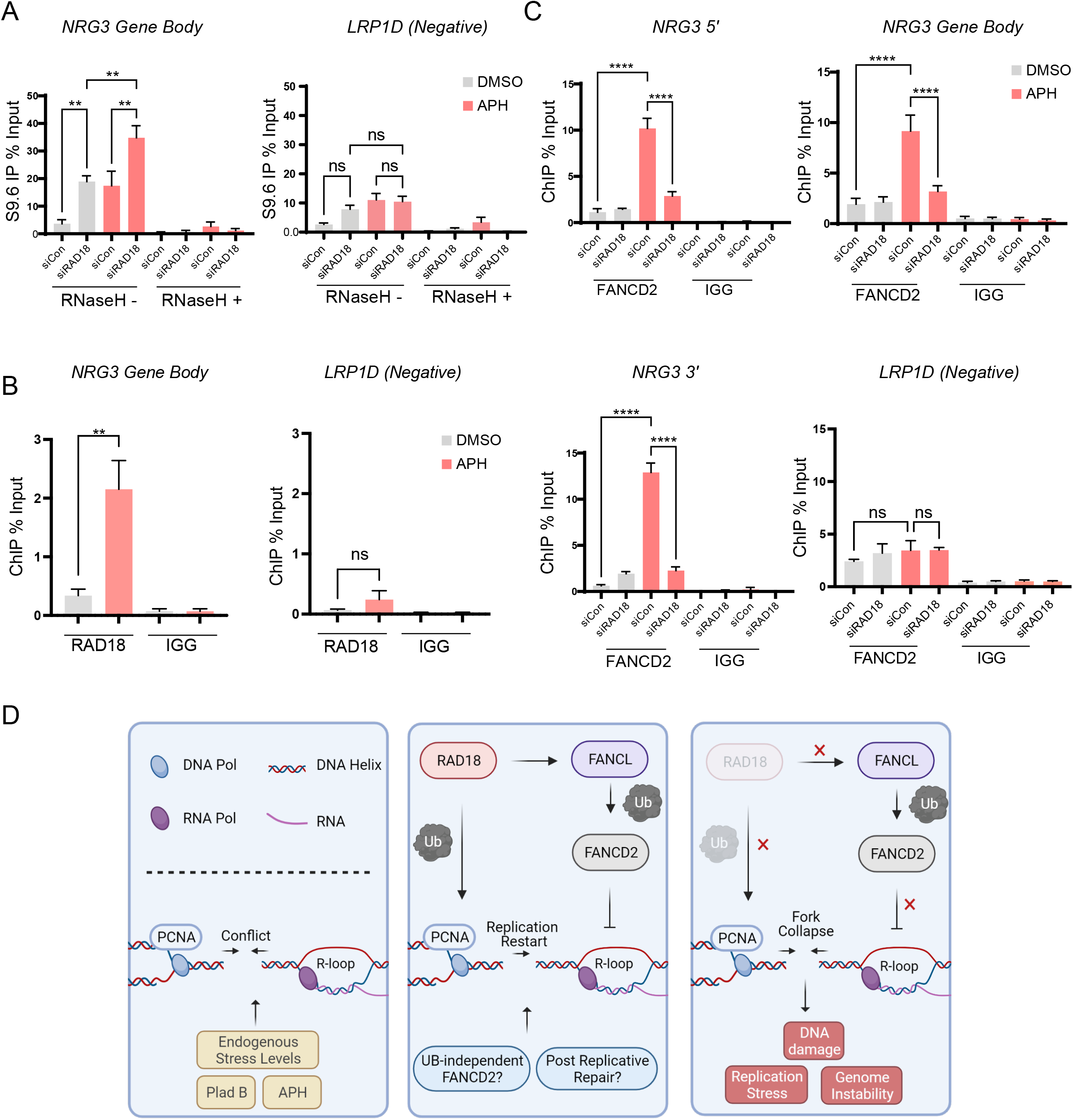
FANCD2 recruitment to Common-Fragile-Sites is dependent on RAD18. (A) DRIP-qPCR in non-targeting control siRNA or siRAD18 treated HeLa cells at common fragile site NRG3 and negative control LRP1D with either a 24 hour treatment of DMSO or APH and with and without an RNaseH1 treatment at 37°C for 48 hours. (B) RAD18 ChIP at common fragile site NRG3 with either a 24 hour treatment of DMSO or APH and IGG ChIP was performed in parallel as a negative control. (C) FANCD2 ChIP in non-targeting control siRNA or siRAD18 HeLa cells at common fragile site NRG3 at 5’, gene body, and 3’ gene locations with LRP1D as a negative control. Cells were treated with either a 24 hour treatment of DMSO or APH and IGG ChIP was performed in parallel as a negative control. (D) Model. Conflicts arising endogenously or due to replication or transcriptional stress can trigger PCNA-Ub (left). Rad18, through its effects on PCNA, FANCD2 recruitment and possibly other pathways, ensures timely resolution of conflicts and associated R-loops (center). In the absence of Rad18, R-loops and conflicts persist, FANCD2 recruitment is reduced, and DNA damage occurs (right). Treatment concentrations of APH (400 nM) and RNaseH1 (5 units). (A-C) ANOVA, p<0.05, n=3. Error bars represent standard deviation. *DRIP*, DNA-RNA immunoprecipitation; *siCon*, non-targeting control siRNA; *APH*, aphidicolin; *Negative*, negative control loci; *ns*, not significant; *, p<0.05; **, p<0.01; ***, p<0.001; ****, p<0.0001.

## Discussion

RAD18, through its effects on PCNA mono-ubiquitination, is a hub for replication fork protection, restart, and lesion bypass (7). The stresses that require RAD18 activity for tolerance are varied and DNA damage can trigger both replication-dependent and independent RAD18 responses (50). Additionally, cells without RAD18 have higher levels of sister-chromatid exchanges (SCEs) even in the absence of external stress (51), suggesting an important role of RAD18 in dealing with endogenous stress. Here we show that transcription and associated R-loop structures are an important endogenous source of replication stress that require RAD18 for tolerance.

RAD18 mediated PCNA ubiquitination has multiple consequences for replication fork maintenance. Fork reversal and template switching are fork rearrangements triggered by ubiquitin ligases HLTF and SHPRH with the E2 ubiquitin conjugating enzyme UBC13 by promoting PCNA polyubiquitination from a mono-ubiquitinated state (52). RAD18 deficient cells are unable to reverse forks, and therefore Primpol activity would lead to replication gaps (53). Gap filling synthesis by TLS polymerases in S or G2 also requires PCNA ubiquitination (48). Another study showed that RAD18 and PCNA ubiquitination activate a Lig1-Atad5-Caf1 pathway of fork protection which opposes degradation by DNA2 (55). In this same study, defective Okazaki fragment processing impaired PCNA unloading by ATAD5 and subsequent chromatin loading by CAF1 (55). We and others previously showed that ATAD5 mediated PCNA unloading is an important anti-R-loop mechanism (29). Therefore, RAD18-deficient cells likely accumulate R-loops due to replication defects, but are also unable to mount an appropriate response to transcription-associated replication stress. Our results showing that RAD18-deficient cells have persistent replication stress, transcription-replication conflicts, and R-loops are consistent with this breakdown of the orchestration of replication fork signaling. Our data also show that a key component of this defect is failure to recruit the FANCD2 protein to sites of transcription-associated replication stress. These findings underscore the importance of transcription as an endogenous source of replication stress. However, it is highly likely that there is complexity built into this recruitment and that FANCD2 has roles in resolving conflicts beyond RAD18 activation. Indeed, non-canonical ubiquitin-independent functions for FANCD2 at TRCs have also been observed (28). R-loop-dependent activation of ATR is dependent upon fork reversal, therefore fork reversal may be stimulated by R-loop-stalled replication forks to limit genome instability, a process that would be considerably disrupted in RAD18-deficient cells (53,54). Furthermore, defects in this canonical PRR pathway result in ssDNA gaps that promote R-loops that form on DNA distinct from replication forks (35). The type of stress, genomic context, and cell cycle stage might all influence which lesions require the RAD18-FANCD2 pathway described here.

Our data highlights an additional role for RAD18 in reducing unresolved transcription-replication conflicts, thereby reducing R-loop levels, replication stress, and DNA damage. To explore the mechanistic connection between RAD18 and transcriptional stress, we investigated the Fanconi Anemia pathway, due to the proposed role of RAD18 in localizing FANCL to chromatin (14). Furthermore, FANCD2 has been proposed to have multiple roles in both R-loop resolution and conflict tolerance (23, 24, 26, 27). This may be specifically relevant at CFSs where, when confronted with conflicts, FANCD2 promotes fork protection, replication progression, and restricts hybrid formation while also regulating dormant origin firing (48,49,56). Our work strengthens the links between RAD18 and conflicts, identifying the importance of RAD18 dependent FANCD2 recruitment to CFSs and other R-loop prone loci. We highlight the distinction between FANCD2 recruitment induced by ICLs compared to R-loops, suggesting that RAD18 plays a much more important role at R-loops and the associated conflicts. This difference may be linked to the structures underlying these repair pathways. Convergent replication forks are important for ICL repair (55), but conflicts are likely resolved by transactions at a single replication fork. The Fanconi Anemia pathway has been well-characterized in ICL repair, however the upstream factors specific for FANCD2 localization for transcription-associated replication stress have not. For instance, both ubiquitin-dependent and -independent roles of FANCD2 have been reported to limit the genome instability caused by transcriptional stress (24, 28). Moreover, while the XPF nuclease is important for ICL unhooking, and has been implicated in cleavage of R-loops (33, 41) our data suggest that there are important differences between ICL and conflict repair. Transcription is an omnipresent challenge for the replication of genomes and our data works to progress our understanding of the mechanisms that prevent transcription-dependent DNA damage and genome instability. The relative extent of how PRR and FA genes impact R-loops is still uncertain, and could depend on the type of collision or presence of co-factors. Further research identifying both the signaling mechanisms upstream of FANCD2 as well as the relevant nucleases or other factors recruited by FANCD2 to promote replication restart after a transcription-replication conflict will improve our understanding of this important endogenous stress response.

## Acknowledgements

P.C.S. is a Canadian Institutes of Health Research (CIHR) New Investigator and Michael Smith Foundation for Health Research Scholar. The work was supported by the Canadian Cancer Society Innovation to Impact grant, and a CIHR project grant to P.C.S. J.P.W. held a CIHR Canada Graduate Scholarship -M, and a Laurel L. Watters Research Fellowship.

## SUPPORTING INFORMATION

**Figure S1.**
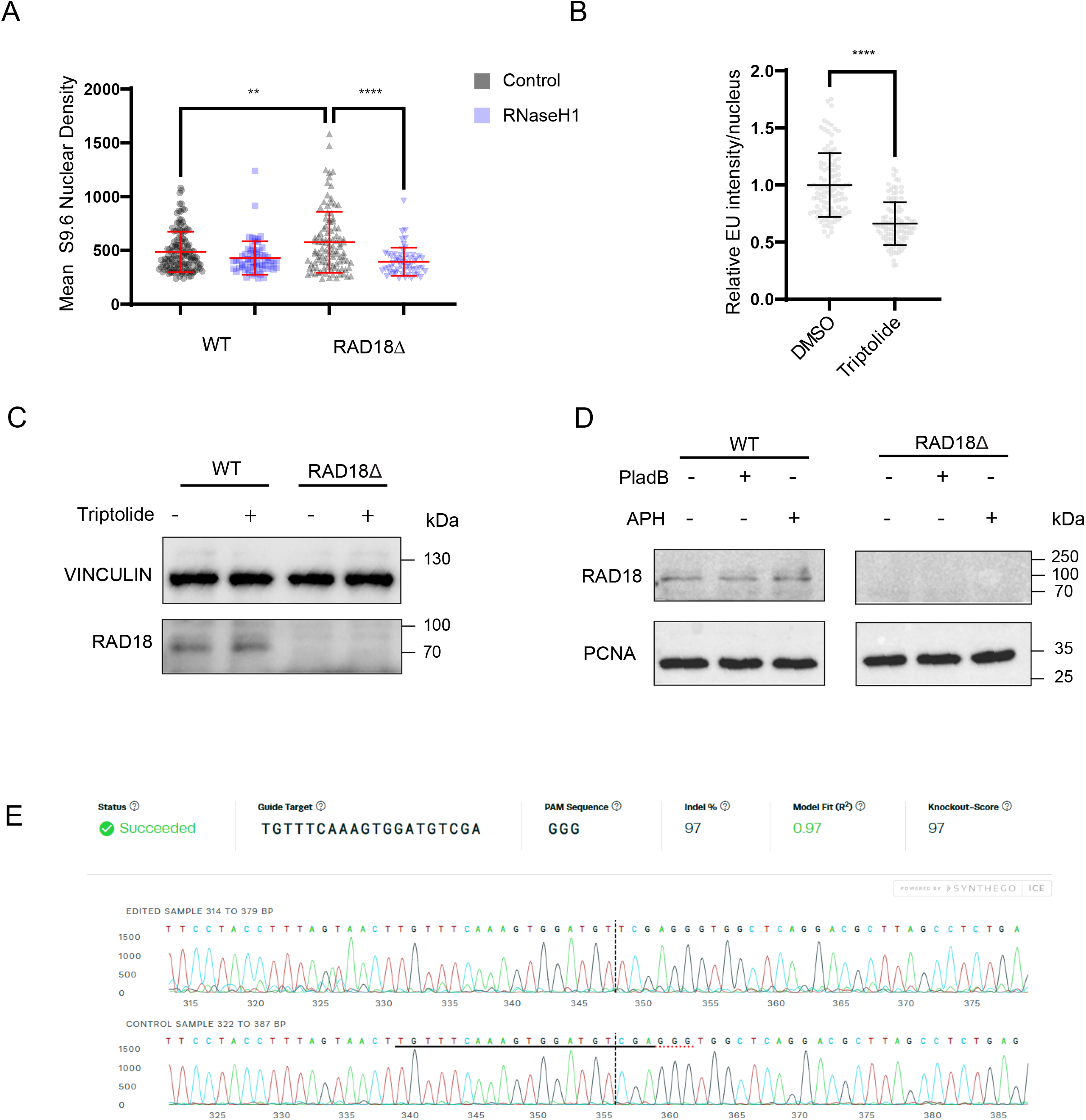
RAD18 CRISPR generated knockout cells have increased DNA:RNA hybrids. (A) S9.6 antibody staining in WT and CRISPR-generated RAD18 knockout clone (KO) of P53^-/-^ RPE1-hTERT cells with or without transfection of a GFP-tagged RNaseH1 overexpression plasmid. (B) Ethynyl uridine (EU) fluorescence measuring active transcription in HeLa cells treated for 2 hours with triptolide. (C) Western blot of P53^-/-^ RPE1-hTERT cell lysate of parental and RAD18 KO cells after a 2 hour triptolide treatment or DMSO. (D) Western blot of P53^-/-^ RPE1-hTERT cell lysate of parental and RAD18 KO cells after a 2 hour Pladienolide B or 24 hour APH treatment compared to a DMSO control. (E) Target site sequences for Crispr/cas9 cleavage were identified using the online synthego CRISPR Design Tool (https://www.synthego.com) and knockout scores were determined for clones and selected using the synthego ICE tool. Treatment concentrations of triptolide (40 nM), Pladienolide B (5 µM), and APH (400 µM). (A) ANOVA p<0.05, N=3. (B) T-test, p<0.05, N=3. Error bars represent standard deviation. *EU*, ethynyl uridine; *PladB*, Pladienolide B; *APH*, aphidicolin; *ns*, not significant; *, p<0.05; **, p<0.01; ***, p<0.001; ****, p<0.0001.

**Figure S2.**
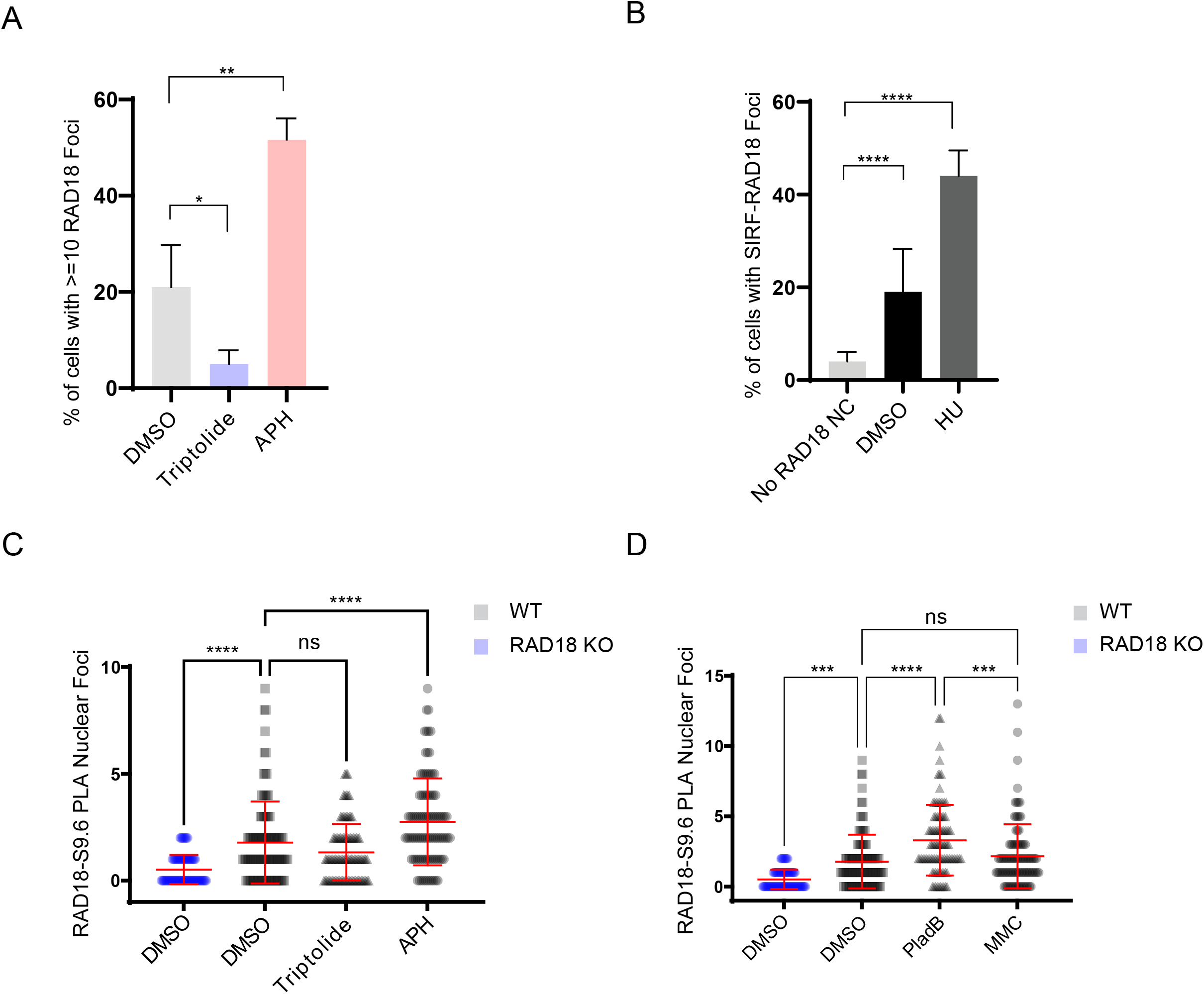
RAD18 localization is dependent on transcription and R-loop inducing stressors. (A) Suppression and stimulation of spontaneous RAD18 foci in triptolide and APH treated HeLa cells for 2 hours and 24 hours respectively. (B) RAD18 localization to nascent DNA replication was performed in HeLa cells after 2 hour treatments of hydroxyurea using the SIRF assay, a PLA experiment bewteen nascent DNA (EDU) and RAD18. (C,D) Suppression and stimulation of RAD18 localization to R-loops in P53^-/-^ RPE1-hTERT WT cells after 2 hr triptolide and 24 hr APH treatment (C) and 2 hour treatments of Pladienolid and Mitomycin C (D). RAD18 KO cells were used as a negative control. Treatment concentrations of triptolide (40 nM), Pladienolide B (5 µM), hydroxyurea (2mM), APH (400 nM), and Mitomycin C (1.5 µM). (A-B) Fisher’s Exact Test. (C-D) ANOVA, p<0.05, n= 3. *SIRF*, Quantitative in situ analysis of protein interactions at DNA replication forks; *PladB*, Pladienolide B; *HU*, hydroxyurea; *APH*, aphidicolin; *MMC*, Mitomycin C; *no RAD18 NC*, negative control lacking the RAD18 antibody; *ns*, not significant; *, p<0.05; **, p<0.01; ***, p<0.001; ****, p<0.0001.

**Figure S3.**
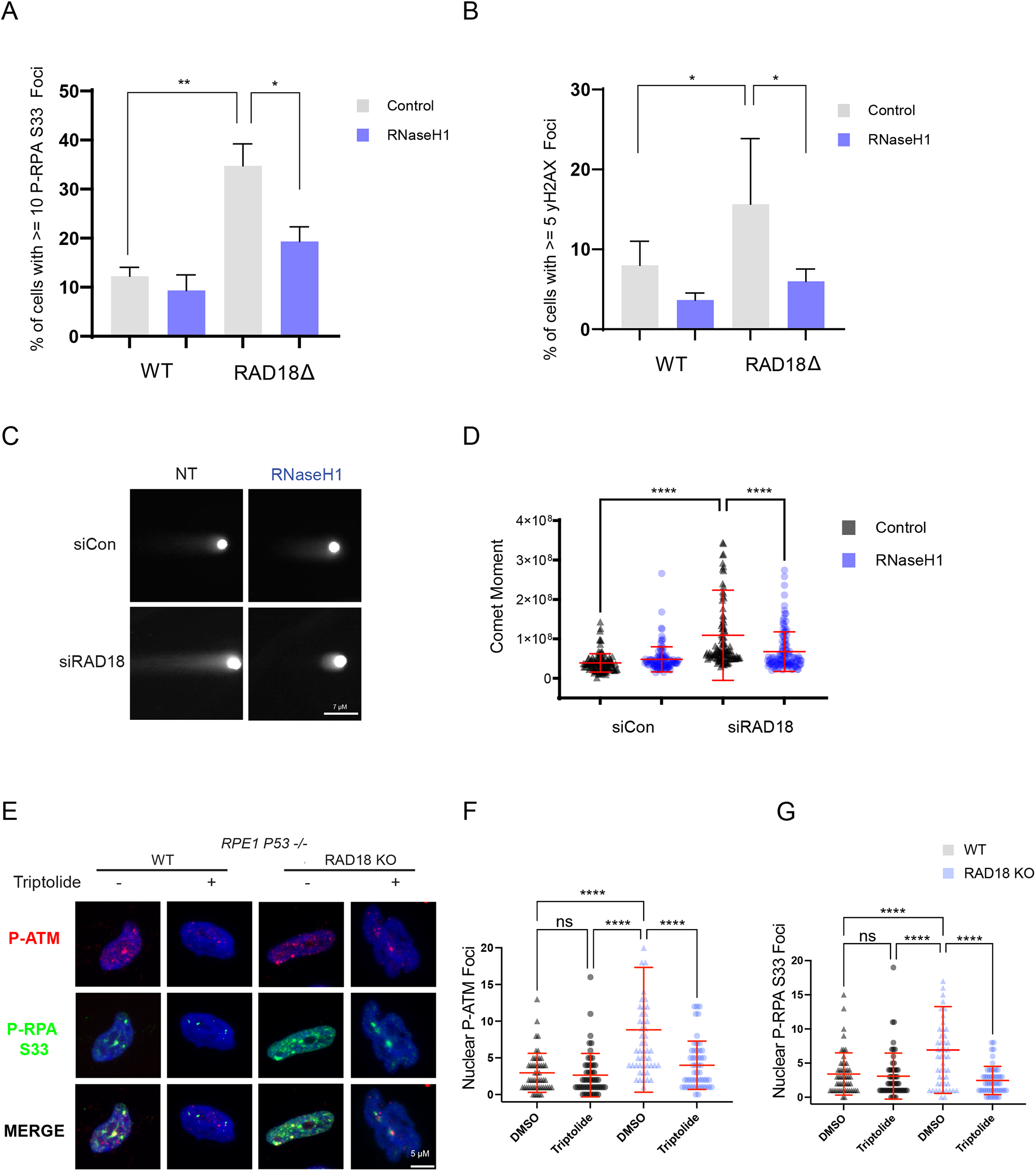
Replication stress in RAD18-KO and transcription-dependent DNA damage. (A) RPA2-ser33 phosphorylation immunofluorescence in P53^-/-^ RPE1-hTERT WT and RAD18 knockout clone cells with or without transfection of a GFP-tagged RNaseH1 overexpression plasmid. (B) H2AX-ser139 phosphorylation immunofluorescence in P53^-/-^ RPE1-hTERT WT and RAD18 knockout clone cells with or without transfection of a GFP-tagged RNaseH1 overexpression plasmid. (C,D) Neutral comet tail moment are shown for non-targeting control siRNA or siRAD18 HeLa cells with or without transfection of a GFP-tagged RNaseH1 overexpression plasmid. C, representative image, D, quantification. (E,F,G) Spontaneous ATM-ser1981 and RPA2-ser33 phosphorylation immunofluorescence in P53^-/-^ RPE1-hTERT RAD18 knockout clone cells and WT with and without a 2 hour treatment of triptolide. A, representative image, B-C, quantification. (A-B) Fisher’s Exact Test. (D, F, and G) ANOVA, p<0.05, n=3 for all. Error bars represent standard deviation. *ns*, not significant; *, p<0.05; **, p<0.01; ***, p<0.001; ****, p<0.0001.

**Figure S4.**
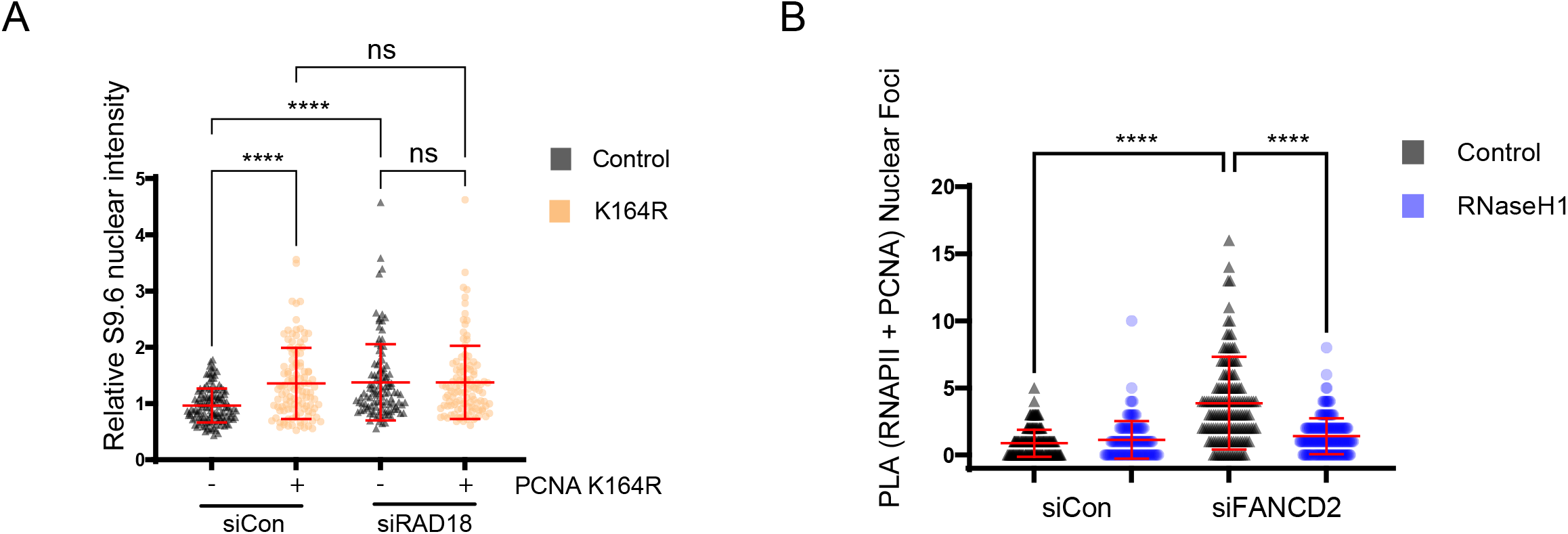
Confirming PCNA-Ub and FANCD2 involvement in R-loops and conflicts. (A) MEF cells carrying a WT (black) or a PCNA^K164R^ (orange) allele were evaluated for S9.6 immunofluorescence with either non-targeting control or siRAD18 treatment. The nucleolus was stained with a nucleolin antibody and S9.6 intensity in these regions was removed to highlight non-nucleolar changes. (B) Proximity ligation of PCNA and RNA polymerase II (RNAP II) in HeLa cells after siRNA depletion of FANCD2 or a non-targeting control, treated with or without a GFP-tagged RNaseH1 overexpression plasmid. (A-B) ANOVA, p<0.05, n=3 for all. Error bars represent standard deviation. *siCon*, non-targeting control siRNA; *ns*, not significant; *, p<0.05; **, p<0.01; ***, p<0.001; ****, p<0.0001.

**Figure S5.**
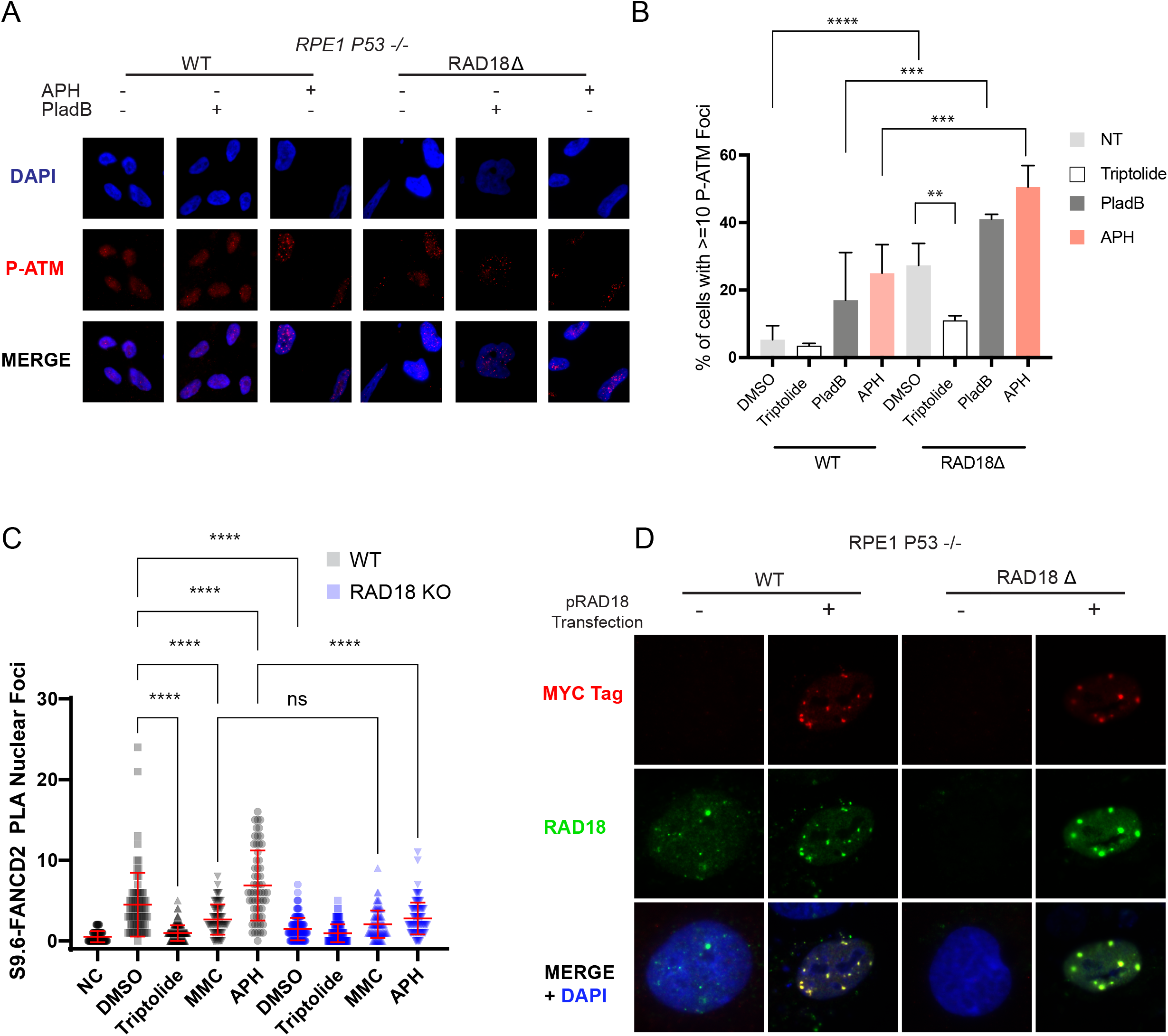
R-loop and conflict inducing stress requires RAD18 for FANCD2 recruitment and limiting DNA damage. (A-B) ATM-ser1981 phosphorylation immunofluorescence in P53^-/-^ RPE1-hTERT RAD18 knockout clone cells and WT with and without a 2 hour treatment of triptolide and pladienolide B and 24 treatment of APH. A, representative image, B, quantification. (C) Stimulation and suppression of FANCD2 localization to R-loops using FANCD2-S9.6 proximity ligation assay after 2 hr treatments of triptolide and Mitomycin C, 24 hour treatments of APH compared to DMSO treatment in P53^-/-^ RPE1-hTERT WT and RAD18 KO cells. (D) Representative image of MYC tag and RAD18 immunofluorescence in P53^-/-^ RPE1-hTERT RAD18 knockout clone cells and WT with and without transfection of MYC-tagged RAD18 CAG plasmid construct. Treatment concentrations of Triptolide (40 nM), Pladienolide B (5 µM), Mitomycin C (1.5 µM) and APH (400 nM). (B,C) ANOVA, p<0.05, n=3 for all. Error bars represent standard deviation. *PLA*, proximity ligation assay; *PladB*, Pladienolide B; *MMC*, Mitomycin C; *APH*, aphidicolin; *NC*, negative control lacking the S9.6 antibody; *ns*, not significant; *, p<0.05; **, p<0.01; ***, p<0.001; ****, p<0.0001.

**Figure S6.**
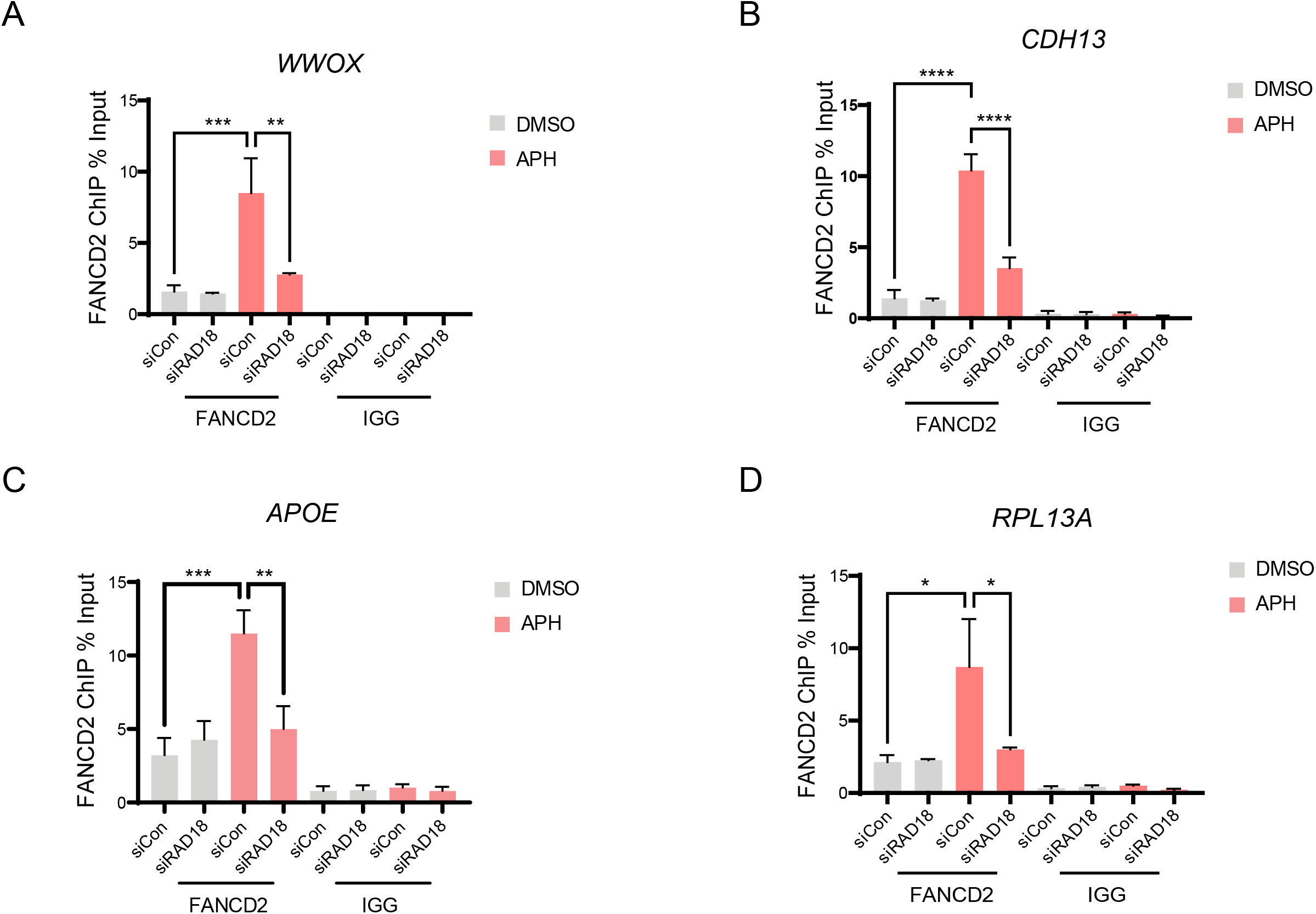
FANCD2 chromatin recruitment to CFS and R-loop prone sites requires RAD18. (A-D) FANCD2 ChIP in Hela cells at common fragile sites WWOX and CDH13 as well as R-loop prone sites APOE and RPL13A. Cells were treated with either a 24 hour treatment of DMSO or APH and IGG ChIP was performed in parallel as a negative control. Treatment concentration of APH (400 µM). ANOVA, p<0.05, n=3 for all. Error bars represent standard deviation. *APH*, aphidicolin; *ns*, not significant; *, p<0.05; **, p<0.01; ***, p<0.001; ****, p<0.0001.

**Table S1.**
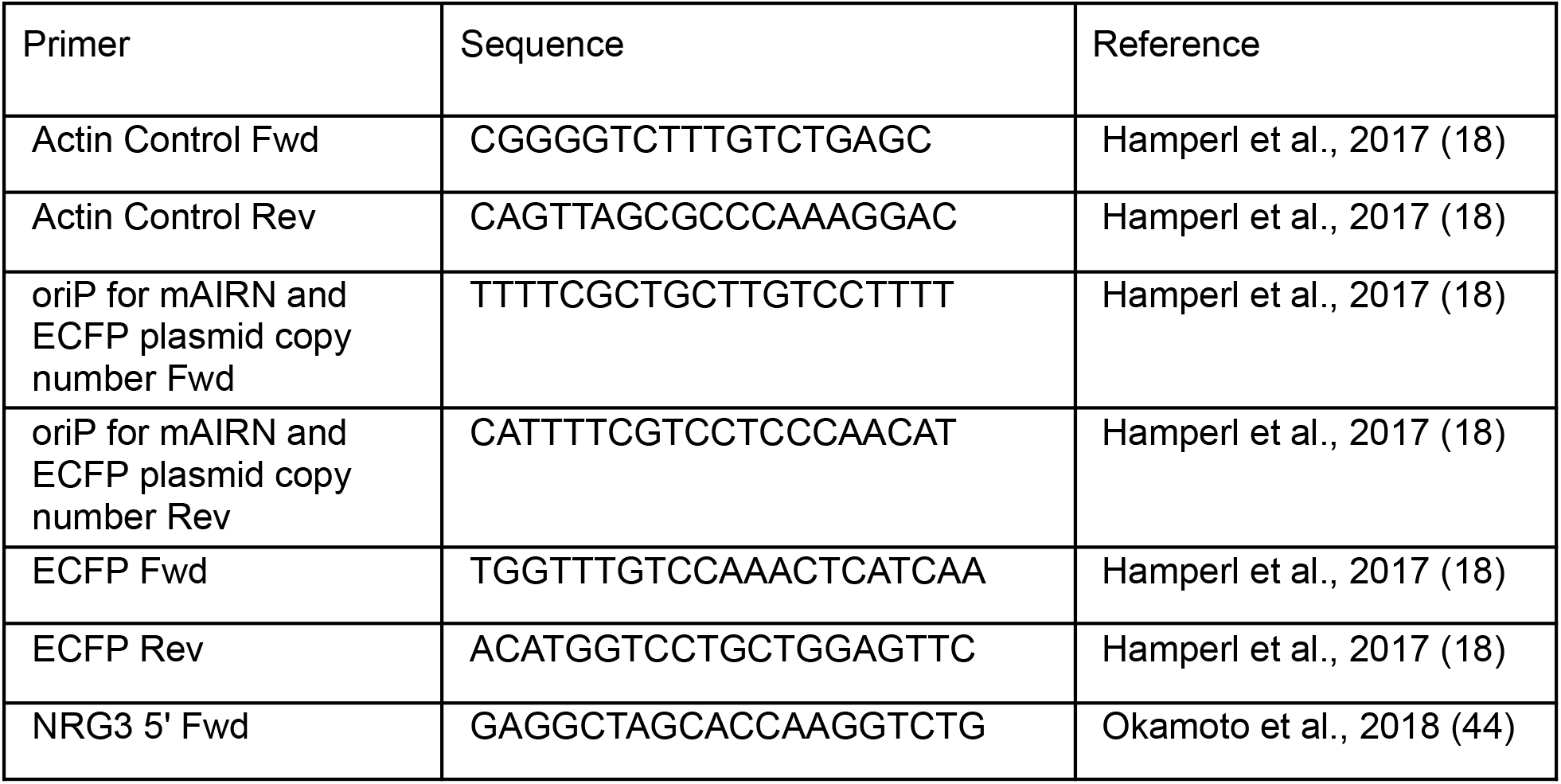
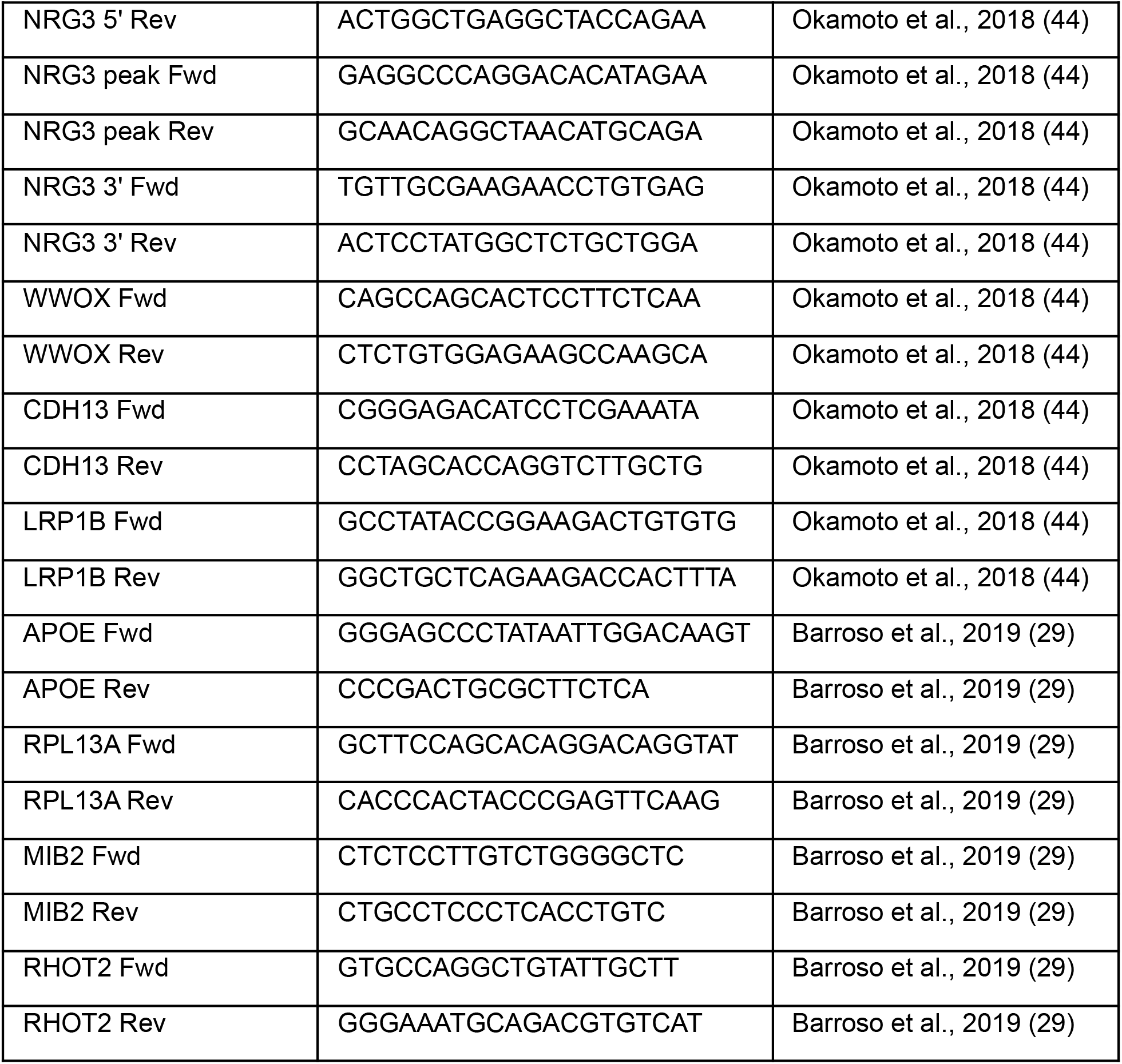
Primer sequences from this study.

**Table S2.**
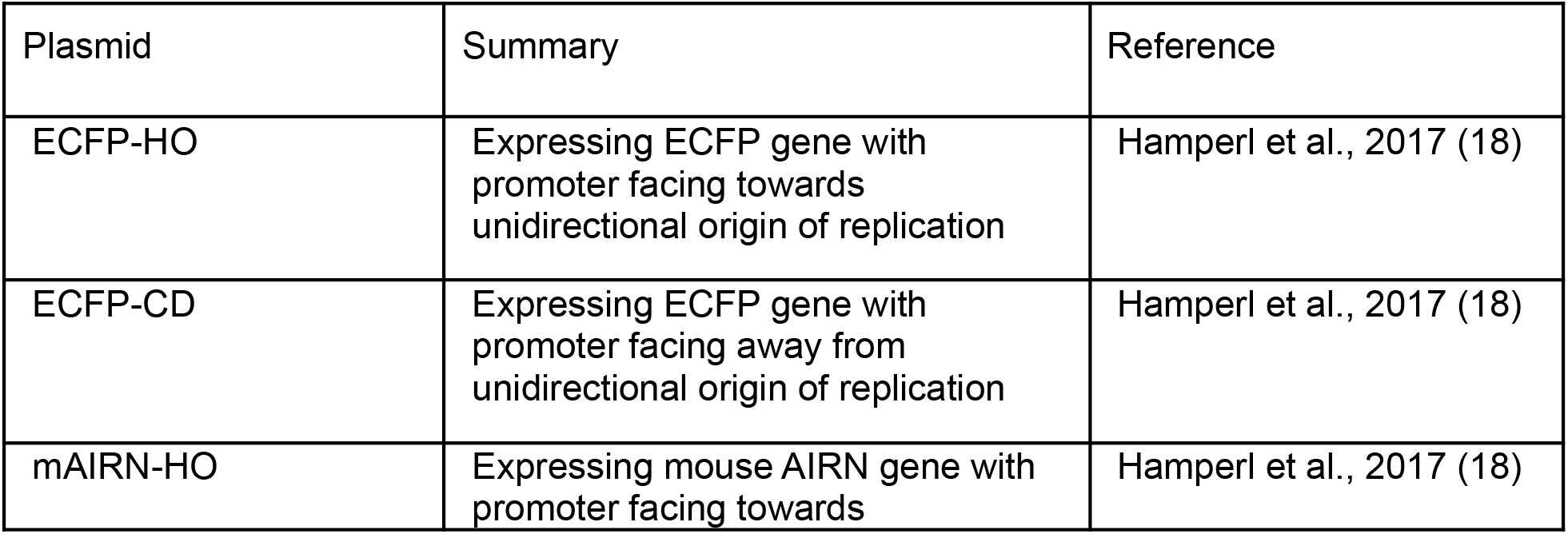
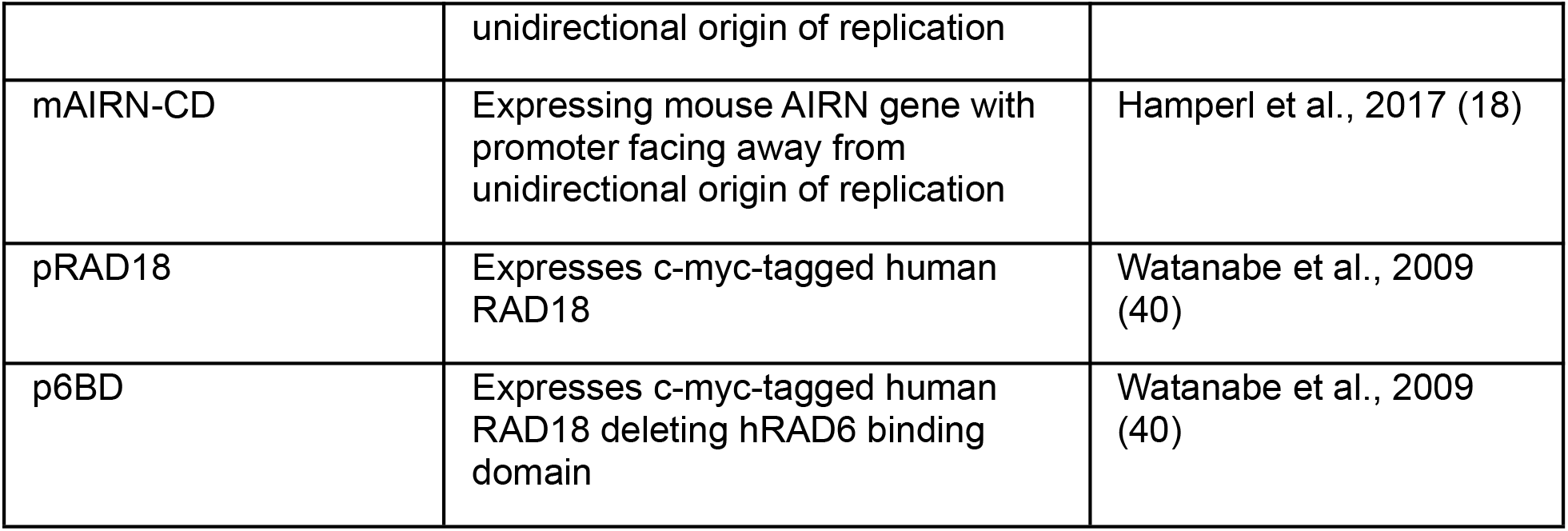
Plasmids from this study.

